# Multivalent assembly of PAR-3 / aPKC complexes establishes cell polarity in *C. elegans* zygotes

**DOI:** 10.1101/2025.04.16.649205

**Authors:** Sheng-Ping Hsu, Daniel J. Dickinson

## Abstract

Cell polarity is essential for the formation and function of animal tissues. Atypical protein kinase C (aPKC), its cofactor PAR-6, and scaffold protein PAR-3 regulate cell polarity in many different animal cell types. PAR-3 oligomerization is important to establish cell polarity, but how oligomerization relates to the assembly of the PAR-3 / aPKC / PAR-6 complex is still unclear. Here, we use *in vivo* and *ex vivo* single molecule techniques to demonstrate cooperativity between PAR-3 oligomerization and its binding to aPKC / PAR-6 in the *C. elegans* zygote. Using genetic perturbations, we demonstrate that aPKC and PAR-6 have independent binding sites for PAR-3. We propose that multivalency drives cooperativity because a single aPKC / PAR-6 heterodimer can interact simultaneously with multiple PAR-3 molecules in an oligomer. Although single binding site mutations do not fully eliminate PAR-3 / aPKC / PAR-6 binding, they do abolish anterior-posterior polarity, demonstrating that PAR-3 / aPKC cooperativity is essential for polarity establishment. Finally, PAR-3 / aPKC cooperativity is downregulated in polarity maintenance, and this downregulation depends on the mitotic kinase PLK-1. Together, our results show how cells can developmentally regulate multivalent assembly of a key polarity complex to achieve timely segregation of cell fate determinants.

## Introduction

Animal tissues require cell polarity for their normal development and function. Cell polarity compartmentalizes the cell into different regions that play different roles. For example, epithelial cells have an apical domain facing the lumen, while the basal domain adheres to the basement membrane [1, 2]. Neurons polarize their neurites to ensure that neural signals can be received by dendrites and transmitted by axons [3–5]. During development, cell polarity maintains stem cell identity, allows differentiation of daughter cells, and contributes to axis specification for whole embryos [6, 7]. Loss of cell polarity interferes with development and disrupts physiology, leading to disease [8, 9]. It is, therefore, of fundamental importance to understand how signaling molecules partition within a cell to establish polarity and how these molecules are regulated by developmental cues.

Atypical protein kinase C (aPKC), its cofactor PAR-6, and the scaffold protein PAR-3 are evolutionarily conserved throughout animals and regulate cell polarity in diverse contexts, including epithelial tissues, neurons, and developing embryos [10–21]. These proteins localize asymmetrically within the cell and dictate polarity by influencing the cytoskeleton, vesicle trafficking, cell junction formation, and cell fate determinants [16, 18, 19, 21–29]. Loss of either PAR-3, PAR-6, or aPKC abolishes polarity, leading to disease and developmental failure [16, 17, 19–21, 26, 29–33]. PAR-3 interacts directly with aPKC/ PAR-6 [17, 34–39] and also binds to phosphoinositides in the plasma membrane [40–43]. PAR-3 is thereby thought to target aPKC to specific locations, and it may also modulate aPKC kinase activity to establish and maintain cell polarity [39, 44–47]. PAR-3 forms homo-oligomers via its N-terminal domain [48, 49], and mutations that block PAR-3 oligomerization abolish cell polarity [45, 50–53]. PAR-3 oligomerization is necessary at least in part because PAR-3 oligomerization increases the avidity of membrane binding, thereby facilitating PAR-3 segregation due to cortical actomyosin flows [54, 55]. However, a previous study also showed that PAR-3 oligomers bind more strongly to aPKC / PAR-6 than monomeric PAR-3 does, implying that PAR-3 oligomerization potentiates aPKC / PAR-6 binding in addition to membrane binding [51]. Yet it remains unclear precisely how PAR-3 oligomerization affects its interaction with aPKC / PAR-6, and how this contributes to polarity.

*In vitro,* PAR-3, aPKC and PAR-6 can interact with one another via multiple binding sites (Figure 1A; [44, 50, 56, 57]). aPKC and PAR-6 interact strongly via heterodimerization of their PB1 domains [34, 36, 57–60]. PAR-3 contains a peptide that is an aPKC substrate (sometimes called conserved region 3, CR3), and this site can mediate stable binding between aPKC and PAR-3 when ATP is absent or aPKC is inactive [44, 61]. In addition, a PDZ ligand at the aPKC C-terminus can bind to the second and third PDZ domains in PAR-3 [39, 56]. PAR-6 has also been reported to bind to the PDZ domains of PAR-3 via its own PDZ domain and PDZ ligand [34, 36, 37, 57], although the significance of this PAR-6 / PAR-3 interaction remains controversial [56]. A challenge in interpreting these *in vitro* experiments is that purification of full-length PAR-3 has not been reported, and full-length PAR-6 is soluble only as a complex with aPKC, so these studies relied on isolated domains rather than full-length PAR-3. As a result, it remains unclear how these different interactions contribute to assembling a PAR-3 / aPKC / PAR-6 complex in cells.

**Figure 1:**
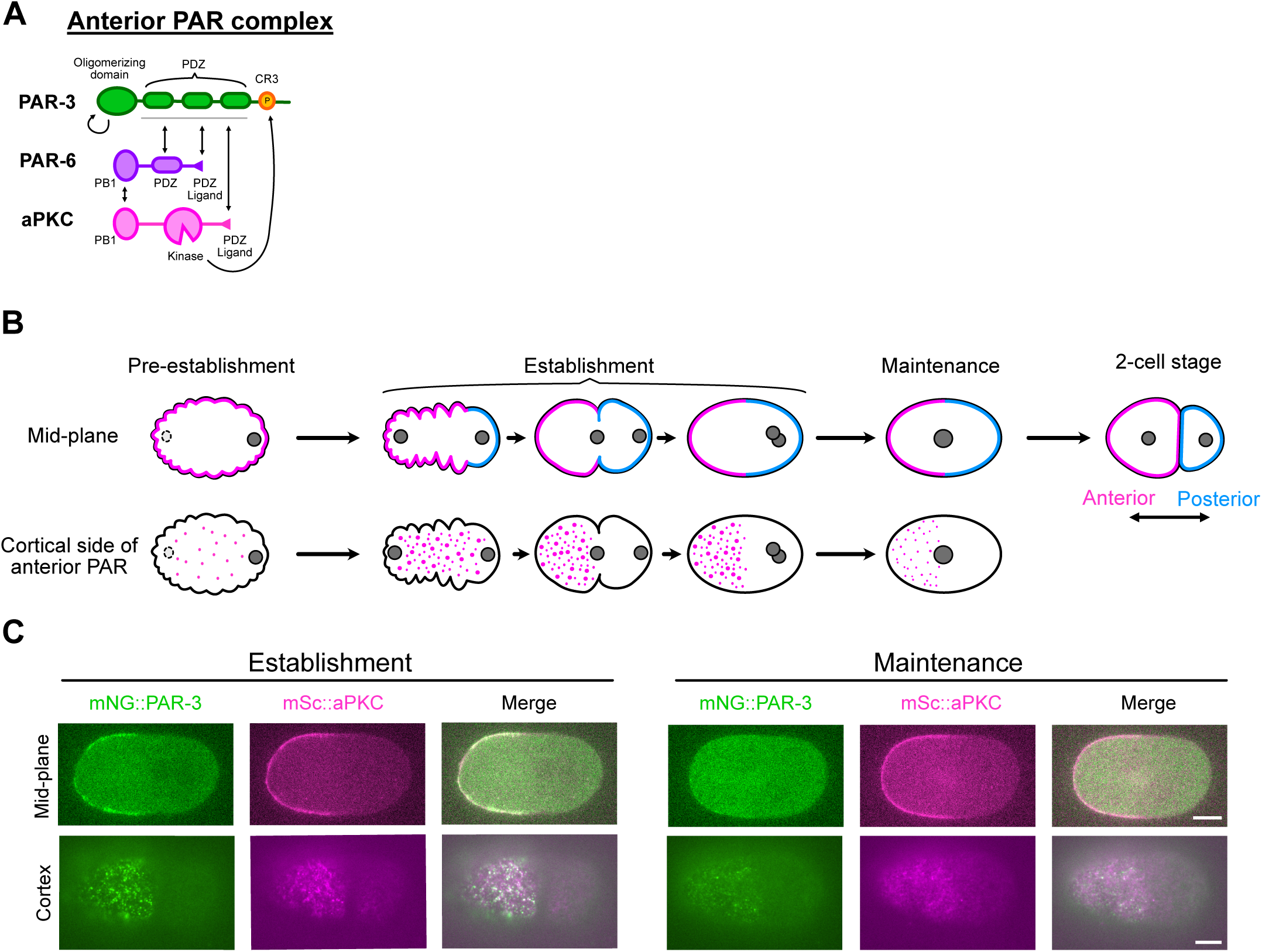
aPAR clustering during polarity establishment in *C. elegans* zygotes. (A) Illustration of the functional domains in PAR-3, PAR-6, and aPKC that interact with each other. (B) Illustration of the morphology and localization of the anterior/posterior PAR complex during the pre-establishment, establishment, and maintenance phase of *C. elegans* zygotes midplane and cortical side. The magenta domain and puncta indicate the anterior PAR proteins, and the blue domain indicates the posterior PAR proteins. (C) Midplane and cortex images of mNG::PAR-3 (green) and mSc::aPKC (magenta) at the indicated stages. Anterior is toward the left. Scale bars represent 10 µm.

Oligomerization of a PAR-3 / aPKC / PAR-6 complex is critical for polarity establishment in the *C. elegans* zygote, which is a well-studied model of cell polarization. in this system, fertilization induces actomyosin contraction, causing anteriorly-directed cortical flows [62, 63] that transport oligomeric clusters of PAR-3, aPKC, and PAR-6 (collectively termed anterior PARs, aPARs) to the anterior [45, 51, 54, 55, 64, 65]. Clearance of aPKC from the posterior cortex allows posterior PAR proteins, including PAR-1 and PAR-2, to occupy the posterior domain, where they antagonize aPARs to maintain anterior-posterior polarity (Figure 1B, [33, 66]). Following the cessation of cortical flows, there is a polarity maintenance phase lasting several minutes, during which opposing pools of aPARs and pPARs are stabilized by a combination of mutual antagonism and positive feedback [33, 66–68]. PAR-3 oligomerization and aPAR clustering are important during both the establishment and maintenance phases. During establishment, PAR-3 oligomerization is critical for coupling of aPAR clusters to cortical flow [51, 54, 55]. During maintenance, PAR-3 oligomerization is reduced due to phosphorylation by the cell cycle kinase PLK-1 [51], but oligomerization still contributes to stable maintenance of the anterior domain (Figure 1B-C [64, 68, 69]). How cells regulate PAR-3 oligomerization during different cell stages is still not fully understood.

Here, we address how PAR-3 oligomerization recruits aPKC / PAR-6 to form the aPAR complex, using single-molecule techniques *ex vivo* and *in vivo*. We show that PAR-3 oligomerization and aPKC / PAR-6 binding are cooperative due to the presence of multiple binding sites that enable multivalent complex assembly. We utilized engineered mutants, drug treatments, and RNAi knockdown to map the key interactions that lead to cooperativity. Finally, we show that PAR-3 / aPKC cooperativity is reduced during polarity maintenance in a PLK-1-dependent fashion, indicating that cooperative aPAR complex assembly is actively regulated during cell polarization.

## Results

### PAR-3 oligomerization cooperates with PAR-3 / aPKC interaction to form aPAR during polarity establishment

To study how PAR-3, aPKC and PAR-6 assemble the aPAR complex, we first performed single-cell, single-molecule pulldown (sc-SiMPull; Figure 2A [51, 58]) from individual establishment-phase *C. elegans* zygotes. We lysed single zygotes carrying endogenously tagged mNeonGreen::PAR-3 along with either HaloTag::aPKC or PAR-6::HaloTag; captured PAR-3 complexes on a coverslip using monovalent anti-mNeonGreen nanobodies; and recorded single-molecule images using TIRF microscopy to visualize binding of labeled complexes to the coverslip in real time (Figure 2A). We used photobleaching step counting of PAR-3 signals to measure the number of PAR-3 molecules in each cluster, while simultaneous landing (termed ‘co-appearance’) of PAR-3 and aPKC / PAR-6 indicated whether each individual PAR-3 oligomer was in complex with aPKC / PAR-6 (Figure 2B [58]). PAR-3 monomers were rarely detected in complex with aPKC or PAR-6, but clusters containing 3 or more PAR-3 molecules were likely to contain aPKC or PAR-6 (Figure 2C). This result is consistent with measurements of the PAR-3 / PAR-6 interaction using an older version of the sc-SiMPull technique [51], and suggests that monomeric PAR-3 has a low binding affinity to aPKC / PAR-6 while PAR-3 oligomers have a higher apparent affinity. The fraction of PAR-3 clusters containing aPKC vs. PAR-6 were indistinguishable, suggesting that aPKC and PAR-6, which interact very strongly with one another [58], are recruited together onto PAR-3 clusters (Figure 2C). Therefore, in the experiments that follow, we used fluorescently tagged aPKC and PAR-6 interchangeably.

**Figure 2:**
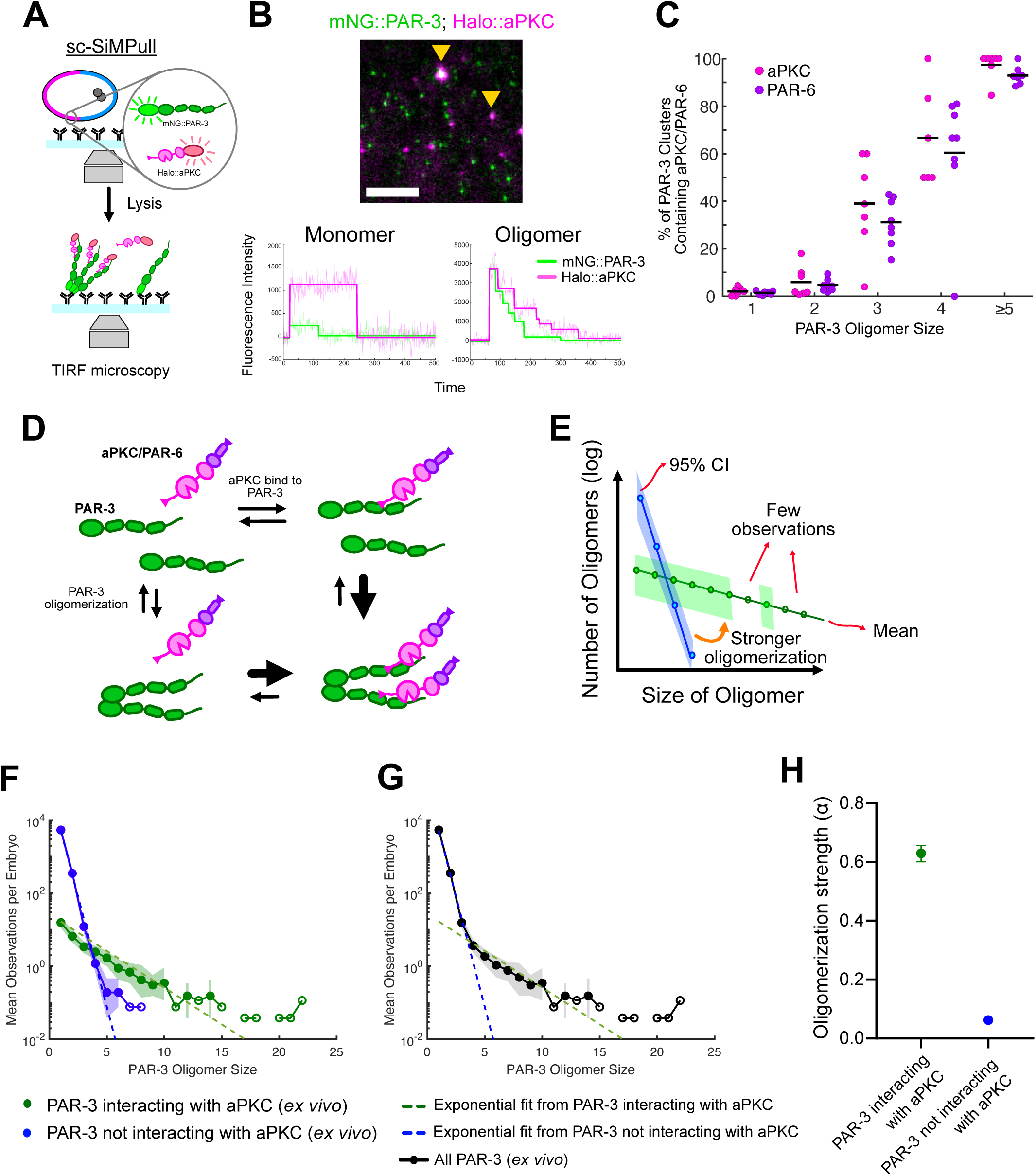
PAR-3 oligomerization and aPKC binding are cooperative during polarity establishment. (A) Illustration of the sc-SiMPull approach. We place a staged embryo into an antibody-coated microfluidic channel whose bottom surface is a coverslip. After lysing the embryo, mNeonGreen::PAR-3 is pulled down on the coverslip surface, and we observe the fluorescent signal on a TIRF microscope. (B) Top: Single-molecule image from sc-SiMPull showing mNG::PAR-3 and Halo::aPKC captured from a single *C. elegans* zygote. The mNG signals co-appearing with the Halo signals are indicated by yellow triangles and oligomeric PAR-3. Scale bar represents 5 µm. Bottom: Fluorescence intensity traces of mNG::PAR-3 and Halo::aPKC classified as monomer (left) and oligomer (right). (C) Fraction of PAR-3 clusters containing aPKC or PAR-6 as a function of size, measured using sc-SiMPull on establishment-phase embryos. n=25,172 PAR-3 complexes from N=7 embryos (aPKC); n=62,296 PAR-3 complexes from N=8 embryos (PAR-6). (D) Illustration of the thermodynamic cycle resulting in assembly of a dimeric PAR-3 / aPKC / PAR-6 complex. (E) Illustration of oligomer size distribution plots. We plot the mean observations per embryo as a function of oligomer size on a semi-log scale. The shaded region shows the bootstrap 95% confidence interval for each dataset. Hollow data points without 95% confidence intervals indicate oligomer sizes that were detected too infrequently to allow the calculation of a meaningful confidence interval (see methods). Linear oligomers are expected to follow an exponential size distribution [68], with a shallower slope indicating stronger oligomerization. (F-H) Comparison of the oligomer size distributions for PAR-3 interacting with aPKC (F, green), PAR-3 not interacting with aPKC (F, blue), and all PAR-3 (G) from sc-SiMPull. Dashed lines are exponential fits to the data (F, G). The oligomerization strength (α) from the exponential fit and its 95% bootstrap confidence interval are plotted in (H). n=149,877 PAR-6 complexes from N=26 embryos.

One possible explanation for the higher aPKC / PAR-6 occupancy of PAR-3 oligomers is that oligomers simply contain more aPKC / PAR-6 binding sites. However, the sharp increase in occupancy from PAR-3 monomers to trimers is incompatible with this hypothesis [51], implying that PAR-3 oligomerization increases aPKC / PAR-6 binding affinity rather than only providing more available binding sites. Moreover, by analyzing photobleaching steps from the HaloTag::aPKC signal, we found an approximately 1:1 ratio of PAR-3 and aPKC molecules in each oligomer, suggesting that all aPKC / PAR-6 binding sites within a PAR-3 oligomer are activated in concert (Supplementary Figure 1A). Although PAR-3 monomers and dimers were infrequently detected in complex with aPKC/ PAR-6, the fraction of molecules bound was still higher compared to a negative control protein, indicating that aPKC / PAR-6 can bind to PAR-3 monomers above background levels (Supplementary Figure 1B). Photobleaching step counting tends to underestimate oligomer sizes due to incomplete fluorophore maturation and simultaneous bleaching of two more fluorophores [51, 70], so we considered the possibility that apparently monomeric PAR-3 / aPKC complexes we detected were in fact larger clusters that were mis-classified as monomers. To test this, we utilized a monomeric PAR-3 mutant lacking the oligomerization domain [55]. The mutated PAR-3 monomer still interacted with aPKC above background levels (Supplementary Figure 1B). This result is consistent with *in vitro* data, which found that (presumably monomeric) PAR-3 fragments lacking the oligomerization domain can bind to aPKC [39, 56]. We conclude that monomeric PAR-3 has a basal affinity for aPKC / PAR-6, which is strongly enhanced in oligomers containing 3 or more subunits.

From a thermodynamic perspective, if PAR-3 oligomerization increases the aPKC / PAR-6 binding affinity of PAR-3 molecules in a cluster, the converse must also be true: aPKC/ PAR-6 binding must promote PAR-3 oligomerization (Figure 2D). To look for such an effect, we analyzed the distributions of PAR-3 oligomer sizes from the same sc-SiMPull data (Figure 2E). We observed that PAR-3 clusters containing aPKC were skewed towards larger sizes compared to clusters not containing aPKC, suggesting that the presence of aPKC / PAR-6 shifts the equilibrium in favor of PAR-3 oligomerization (Figure 2F-H). We conclude that PAR-3 oligomerization and PAR-3 / aPKC binding are cooperative, since these events promote one another (Figure 2D).

PAR-3 oligomerization is thought to occur via stacking of N-terminal domain monomers in a head-to-tail fashion [50]. Since the resulting oligomers are linear, the probability of adding another monomer will not depend on size, and therefore the distribution of oligomer sizes is expected to be exponential with the form *A_n_* = *A*_1_*α^n^*^−1^ where *A_n_* is the abundance of oligomers with n subunits, and 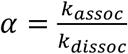 is a parameter that describes the strength of oligomerization [68]. Indeed, both populations of PAR-3 (those with and without aPKC / PAR-6) were well fit by single exponential distributions, indicating that PAR-3 forms linear oligomers in both cases (Figure 2F-G). However, clusters containing aPKC / PAR-6 displayed a distribution with a shallower slope (larger α), indicating stronger oligomerization (Figure 2H). The combined size distribution of all PAR-3 clusters was biphasic, with an inflection point between 3 and 4 monomers per cluster, suggesting a threshold size above which PAR-3 / aPKC / PAR-6 complex assembly is potentiated. Strikingly, this threshold size corresponds approximately to the minimum cluster size (3 monomers) that we previously showed was necessary and sufficient to support polarity establishment [54]. Below, we use this biphasic distribution shape as a signature of cooperative PAR-3 / aPKC / PAR-6 complex assembly.

We next sought to verify these findings from our *ex vivo* sc-SiMPull assay in live cells. To this end, we performed cortical imaging with chimeric labeling (Figure 3A) [71]. Briefly, we utilized *C. elegans* zygotes carrying endogenously tagged mNG::aPKC and Halo::PAR-3. We labeled Halo::PAR-3 with a combination of two HaloTag ligand dyes: a high concentration of far-red dye to label the majority of the molecules, and a lower concentration of a red dye that creates sparse single-molecule labels. The sparse signal facilitates accurate particle tracking and serves as an internal standard for fluorescence intensity calibration [71]. In this way, we measured the size distributions of cortical PAR-3 clusters. By simultaneously imaging mNG::aPKC in the green channel, we determined whether each PAR-3 cluster contained mNG::aPKC (Figure 3B). We plotted the distribution of PAR-3 cluster sizes from this cortical imaging experiment and observed that PAR-3 clusters colocalizing with aPKC had larger sizes than those not colocalized with aPKC, suggesting that aPKC binding enhanced PAR-3 oligomerization *in vivo* (Figure 3C). Compared to the *ex vivo* sc-SiMPull data, the *in vivo* cortical imaging experiment showed a similar size distribution for PAR-3 / aPKC complexes. However, in cortical images, we detected fewer PAR-3 clusters lacking aPKC than in sc-SiMPull experiments (Figure 2F, 3C). This is mostly likely because sc-SiMPull captured all PAR-3 from the cytoplasm and cortex, while *in vivo* cortical imaging only observed membrane-bound PAR-3. We infer that the cell contains a large cytoplasmic pool of PAR-3 monomers and dimers that are not in complex with aPKC, while the cortical pool comprises larger clusters of PAR-3, aPKC and PAR-6. Together, these data reveal cooperative assembly of PAR-3 / aPKC / PAR-6 complexes at the cell cortex during polarity establishment.

**Figure 3:**
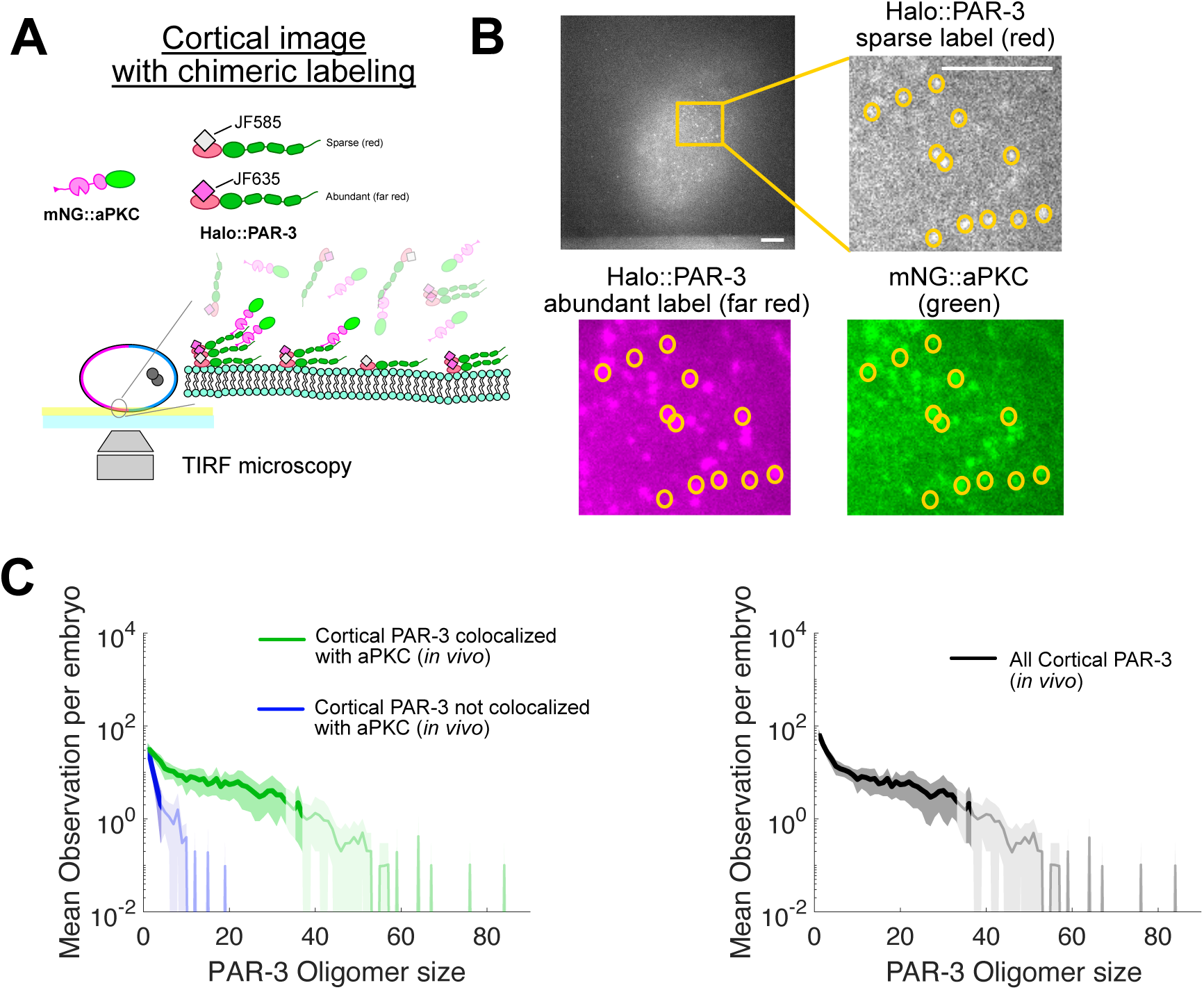
Cortical PAR-3 complexes containing aPKC have stronger oligomerization *in vivo*. (A) Illustration of cortical imaging with chimeric labeling. Staged embryos carrying endogenously Halo-tagged PAR-3 are labeled with two different HaloTag ligand dyes: sparse JF585 and abundant JF635. Cortical PAR-3 clusters are observed using TIRF microscopy. (B) Images acquired from cortical imaging with chimeric labeling. Scale bars represent 5 µm. (C) Comparison of oligomer size distributions for PAR-3 colocalized with aPKC (left, green), PAR-3 not colocalized with aPKC (left, blue), and all PAR-3 (right) measured using cortical imaging with chimeric labeling. The line graph shows the mean and 95% bootstrap confidence interval. The curve is truncated where it passes below 10^-2^; for these complex sizes, there were 0 observations in our dataset. The faint region indicates <20 observations of a given size. n=3,099 PAR-3 clusters from N=10 embryos.

### PDZ/ligand binding is essential for PAR-3 / aPKC cooperativity

We next asked which binding sites drive cooperative assembly of the PAR-3 / aPKC / PAR-6 complex *in vivo.* We first considered the PDZ ligand at the C-terminus of aPKC, which can bind to the second and third PDZ domains of PAR-3 [39, 56]. Mutation at the C-terminal valine of *Drosophila* aPKC disrupted the PAR-3 / aPKC interaction *in vitro* and abolished apical polarization of aPKC in neuroblasts [39, 56]. We mutated this residue (V597 in *C. elegans* aPKC) to alanine and used sc-SiMPull to assess aPAR complex assembly (Figure 4A). Importantly, the binding affinity between aPKC and PAR-6 was unchanged in the aPKC(V597A) mutant compared to WT (Supplementary Figure 2A-B), indicating that the aPKC / PAR-6 heterodimer remained intact. However, PAR-3 oligomerization was strongly decreased in aPKC(V597A) (Figure 4B). The biphasic distribution of PAR-3 oligomer sizes was lost, and the size distribution now closely matched the single exponential distribution predicted for simple oligomerization without cooperativity (Figure 4B, [68]). This was mainly due to reduced sizes of PAR-3 clusters that contained aPKC (Figure 4C, E); the abundance of PAR-3 oligomers lacking aPKC was unchanged compared to wild-type (Figure 4D, E). These data show that the aPKC C-terminal PDZ-ligand binding is essential for PAR-3 / aPKC cooperativity. Strikingly, however, the interaction between PAR-3 monomers/dimers and aPKC was almost unaffected in the aPKC(V597A) mutant (Figure 4C), suggesting that the aPKC(V597A) mutant disrupted cooperativity without entirely abolishing the PAR-3 / aPKC interaction. This implies that one or more additional binding sites can mediate PAR-3 / aPKC / PAR-6 interactions. Nevertheless, the aPKC(V597A) mutation profoundly disrupted zygote polarity: PAR-3 and PAR-6 lost their enrichment at the anterior cortex (Figure 4E). We could still detect a few PAR-3 puncta at the membrane in cortical images, especially during maintenance phase (Supplementary Figure 2F), but these clusters did not flow towards the anterior (Supplementary Figure 2G). The delocalization of aPAR led to symmetric cell division (Figure 4F, G) and 100% embryonic lethality, demonstrating that the PDZ / PDZ-ligand binding in PAR-3 / aPKC is necessary for asymmetric cell division. We conclude that cooperative assembly of PAR-3 / aPKC complexes is essential for *C. elegans* zygote to establish cell polarity during development.

**Figure 4:**
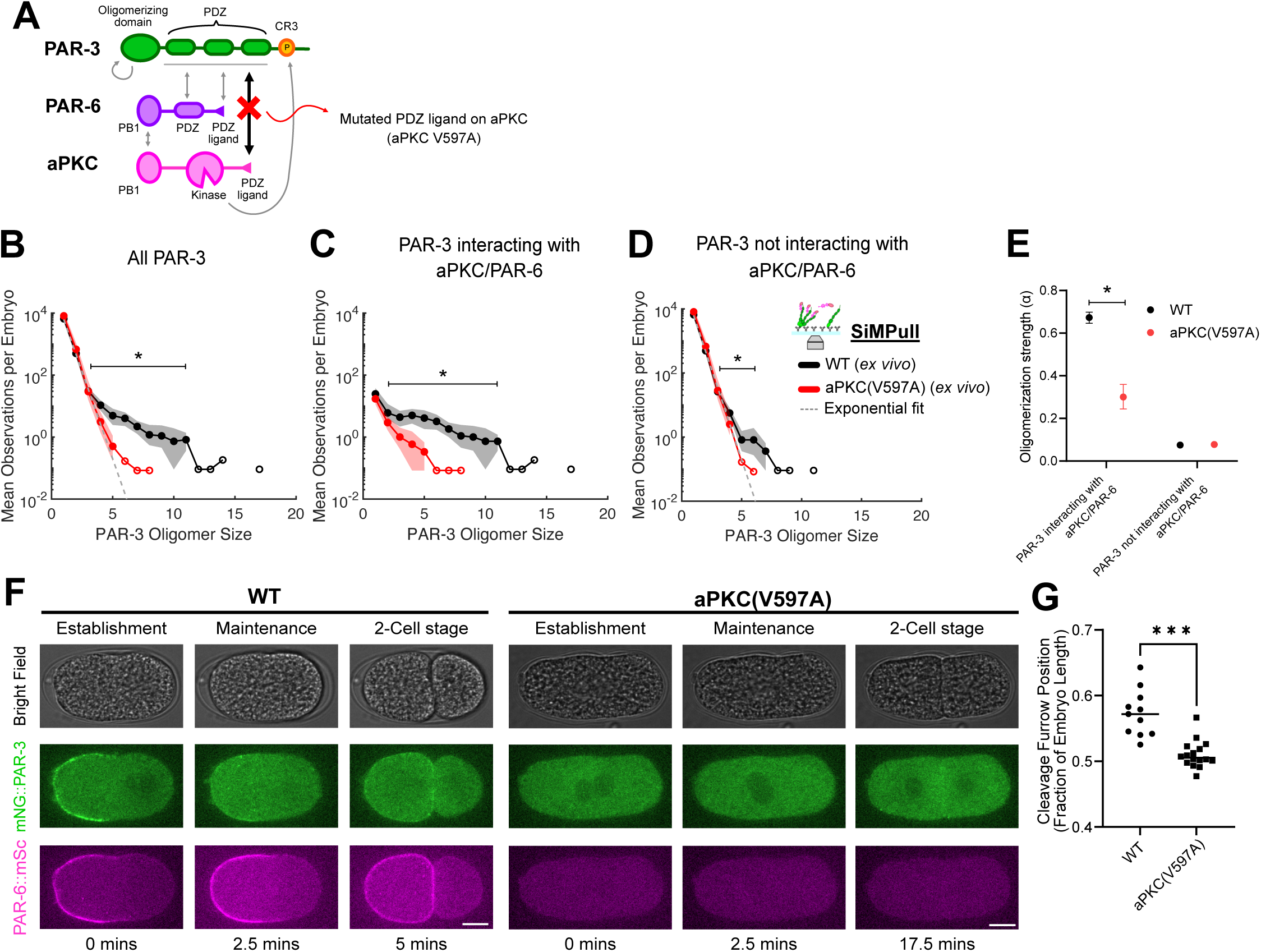
PDZ-ligand mutant of aPKC disrupts cooperativity and abolishes polarity. (A) Illustration of known aPAR complex interactions, indicating the interaction disrupted by the aPKC(V597A) mutant (B-E) Comparison of the oligomer size distributions for all PAR-3 (B), PAR-3 interacting with aPKC / PAR-6 (C), and PAR-3 not interacting with aPKC / PAR-6 (D) measured using sc-SiMPull. Labeling conventions for these plots are illustrated in Figure 2D. * indicates significant differences between groups (based on effect sizes plotted in Supplementary Figure 2C-E). The dashed line is an exponential fit to the population of PAR-3 not interacting with aPKC / PAR-6 in wild-type. (E) The oligomerization strength (α) from the exponential fit and its bootstrap 95% confidence interval. * indicates significant differences between groups because their confidence intervals do not overlap. n=78,185 PAR-3 complexes from N=11 embryos (WT); n=105,830 PAR-3 complexes from N=12 embryos (aPKC(V597)). (F) Midplane images of wild-type and aPKC(V597A) at the indicated stages and carrying the indicated endogenous tags. Anterior is toward the left. Scale bars represent 10 µm. Times are relative to pronuclear meeting, which coincides with the end of the establishment phase. Maintenance phase in wild-type is shown at pronuclear centration, and we show the same interval after the pronuclear meeting for the aPKC(V597A). (G) Length of the AB cell as a fraction of total embryo length in wild-type and aPKC(V597A) 2-cell embryos. **** indicates a significant difference between WT and aPKC(V597A) by Student’s t-test. N=11 embryos (WT); N=16 embryos (aPKC (V597A)).

### PAR-6 contributes to cooperativity in the aPAR complex

PAR-6, as an aPKC cofactor, plays an essential role in forming the aPAR complex [34–36, 41, 56–60, 65, 72–75]. Besides aPKC / PAR-6 interaction, previous studies also provide evidence of a direct PAR-3 / PAR-6 interaction [34–37], which could contribute to assembling the oligomeric PAR-3 / aPKC / PAR-6 complexes we observed (Figure 5A). To determine if PAR-6 is important for PAR-3 / aPKC cooperativity, we knocked down PAR-6 by RNAi and analyzed PAR-3 / aPKC complexes using sc-SiMPull (Supplementary Figure 3A). Similar to the aPKC(V597A) mutant, PAR-6 depletion reduced the sizes of PAR-3 oligomers, especially those in complexes with aPKC (Figure 5B-E). The distribution of PAR-3 oligomer sizes was shifted to a single exponential (Figure 5B), suggesting that PAR-6 is necessary for PAR-3 / aPKC positive cooperativity. However, PAR-6 depletion did not abolish the interaction between PAR-3 monomers and aPKC, which is not surprising considering that the C-terminal PDZ ligand on aPKC remains intact. Thus, PAR-6 contributes to cooperative aPAR complex assembly but is not solely responsible for binding between PAR-3 and aPKC / PAR-6.

**Figure 5:**
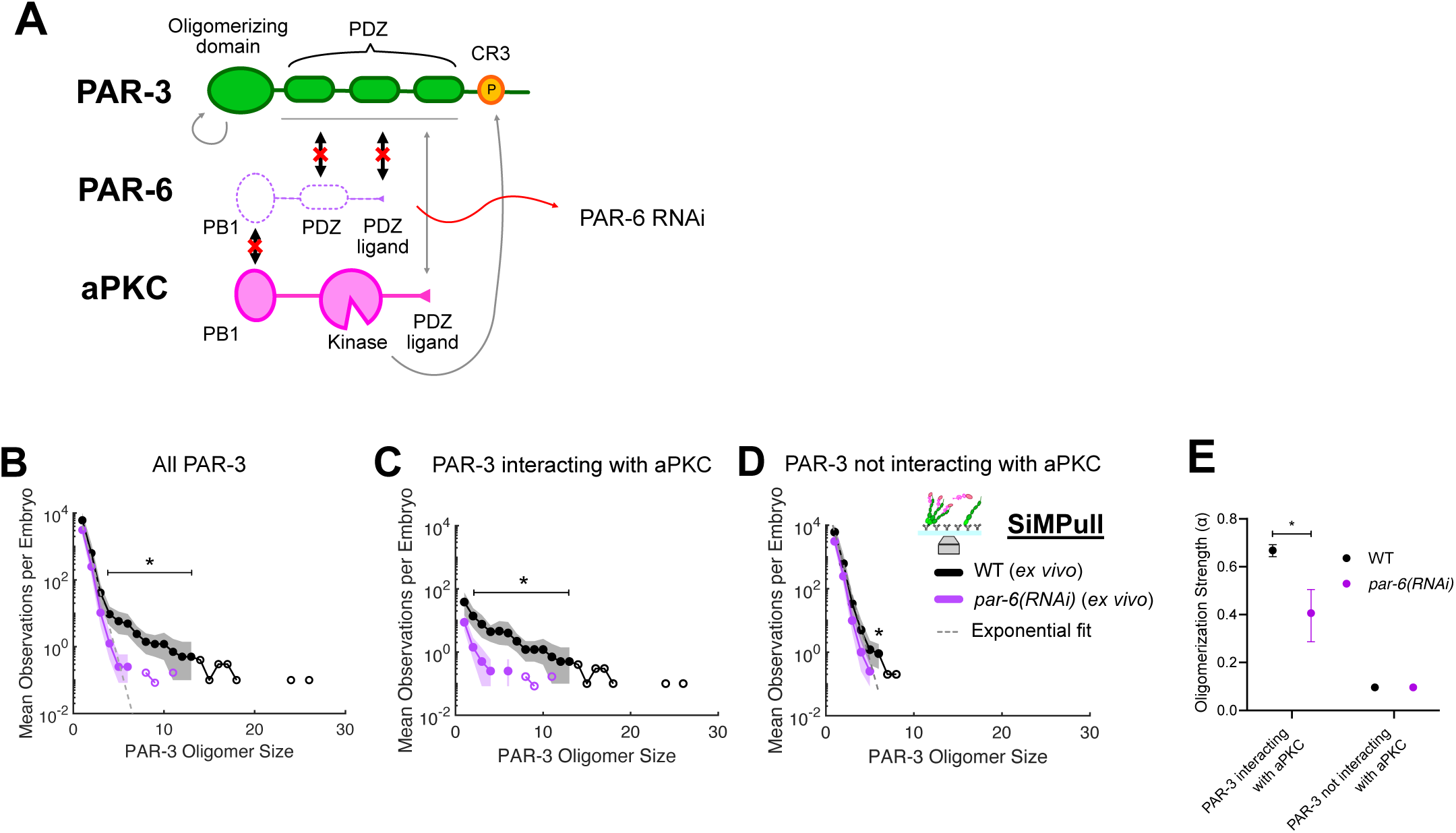
PAR-6 promotes the cooperativity of PAR-3 / aPKC. (A) Illustration of known aPAR complex interactions, indicating the interactions disrupted by *par-6* RNAi. (B-E) Comparison of the oligomer size distributions for all PAR-3 (B), PAR-3 interacting with aPKC (C), and PAR-3 not interacting with aPKC (D) measured using sc-SiMPull. Labeling conventions for these plots are illustrated in Figure 2D. * indicates significant differences between WT and *par-6(RNAi)* (based on effect sizes plotted in Supplementary Figure 3A-C). The dashed line is an exponential fit from PAR-3 not interacting with aPKC in wild-type. (E) The oligomerization strength (α) from the exponential fit and its bootstrap 95% confidence interval. * indicates significant differences between groups because their confidence intervals do not overlap. n= 67,479 PAR-3 complexes from N=10 embryos (WT); n=40,279 PAR-3 complexes from N=12 embryos (*par-6(RNAi)*).

### Kinase-substrate binding of aPKC and PAR-3 is not essential for cooperativity

The aPKC kinase domain can bind stably to its substrate on PAR-3 when ATP is absent [39, 44], but whether this interaction contributes to stable PAR-3 / aPKC interactions in cells remains to be determined. To examine if this kinase-substrate binding can contribute to the PAR-3 / aPKC cooperativity (Figure 6A), we added ATP to the buffer in sc-SiMPull experiments to allow aPKC to complete its catalytic cycle, which should dissolve the kinase-substrate binding of PAR-3 and aPKC as previously shown *in vitro* [39]. ATP addition appeared to slightly shift the distribution of PAR-3 oligomer sizes relative to control conditions, but the difference was smaller than aPKC(V597A) or PAR-6 KD (Figure 6B-E). PAR-3 clusters still exhibited a biphasic size distribution following ATP application (Figure 5B) and the size distributions of complexes containing vs lacking aPKC were still clearly different (Figure 6C-E), suggesting that PAR-3 / aPKC cooperativity still exists under these conditions.

**Figure 6:**
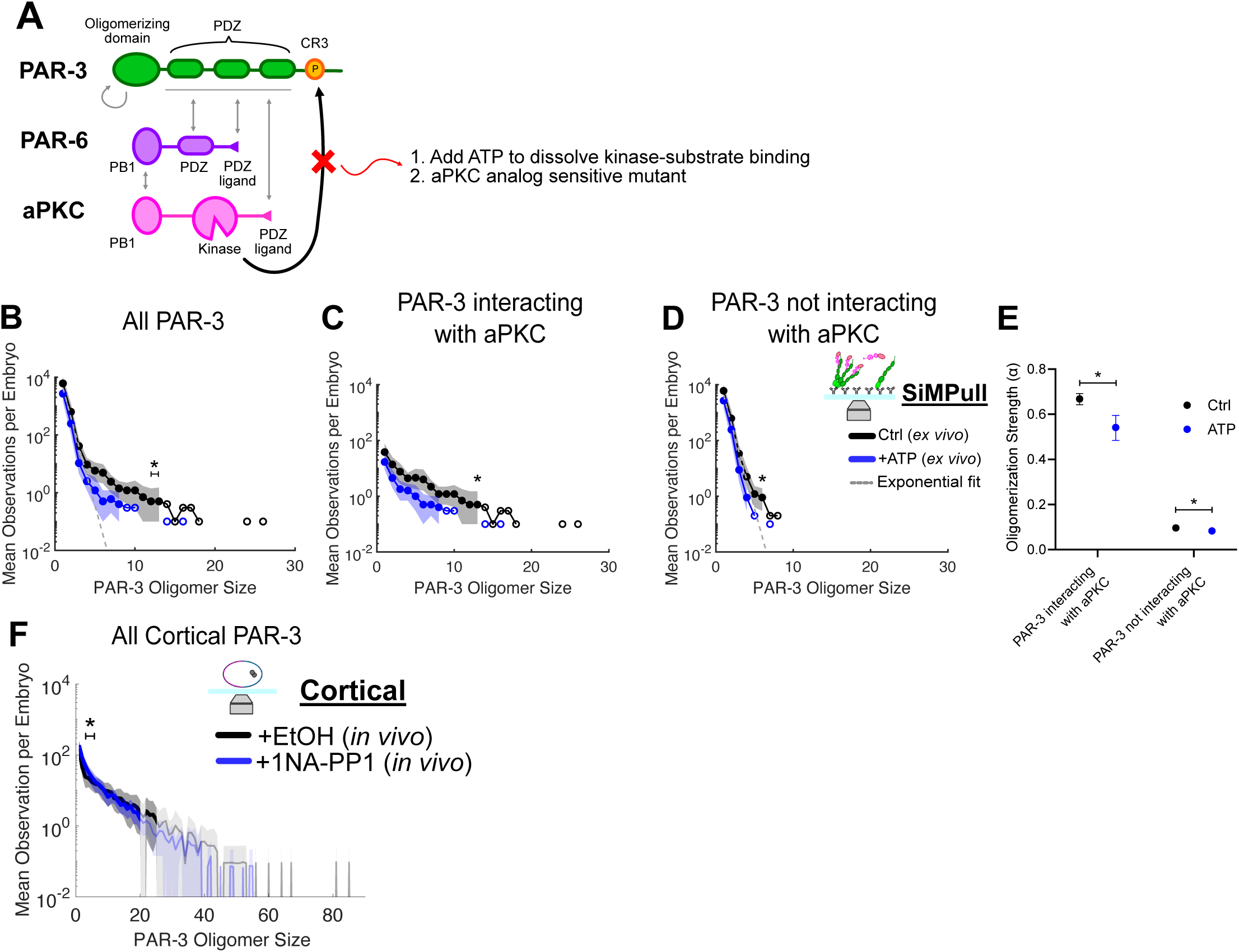
aPKC kinase activity minimally affects the equilibrium of PAR-3 oligomerization *ex vivo* and *in vivo*. (A) Illustration of known aPAR complex interactions, indicating the phosphorylation and kinase/substrate interactions disrupted by our experimental perturbations. (B-E) Comparison of the oligomer size distributions for all PAR-3 (B), PAR-3 interacting with aPKC (C), and PAR-3 not interacting with aPKC (D) measured using sc-SiMPull in the indicated buffer conditions. Labeling conventions for these plots are illustrated in Figure 2D. * indicates significant differences between groups (based on effect sizes plotted in Supplementary Figure 4A-C). The dashed line is the exponential fit from PAR-3 not interacting with aPKC in ctrl. The control dataset (black) is the same data as plotted in Fig. 5B-D and is shown again here for clarity. (E) The oligomerization strength (α) from the exponential fit and its 95% bootstrap confidence interval. * indicates significant differences between groups because their confidence intervals do not overlap. n=67,479 PAR-3 complexes from N=10 embryos (control); n=29,471 PAR-3 complexes from N=10 embryos (+ATP). (F) Comparison of the oligomer size distributions for all PAR-3, PAR-3 colocalized with aPKC, and PAR-3 not colocalized with aPKC from cortical imaging with chimeric labeling in a strain carrying an aPKC analog-sensitive allele and treated with ethanol control or 1NA-PP1. The line graph shows the mean and its bootstrap 95% confidence interval. * indicates significant differences between groups (based on effect sizes plotted in Supplementary Figure 4D). The curve is truncated where it passes below 10^-2^; for these complex sizes, there were 0 observations in our dataset. The faint region indicates <20 total observations of a given size. n=4,175 molecules from N=11 embryos (EtOH); n=7,027 molecules from N=14 embryos (1NA-PP1).

To further test whether aPKC kinase activity influences PAR-3 / aPKC cooperativity *in vivo*, we utilized a *C. elegans* strain containing an analog-sensitive aPKC mutant (aPKC- as) [76] and performed cortical image with chimeric labeling. In this mutant, aPKC-as has normal kinase activity and can support polarity establishment in the ethanol (EtOH) control, but its kinase activity is inhibited by applying the ATP analog, 1NA-PP1 (Supplementary Figure 4E-G). If stable kinase-substrate binding contributed to PAR-3 / aPKC cluster assembly, then we would expect to see larger PAR-3 clusters upon 1NA-PP1 treatment, because 1NA-PP1 would block aPKC from completing its catalytic cycle and releasing phosphorylated PAR-3. However, we observed the opposite: the size distribution of cortical PAR-3 clusters was shifted slightly smaller in 1NA-PP1, mainly due to the presence of more cortical PAR-3 dimers and trimers in the 1NA-PP1 condition (Figure 5E). Moreover, there was no significant change in the abundance of larger PAR-3 oligomers (Figure 6E, Supplementary Figure 4H). Of note, we found that cortical flow was downregulated when we applied 1NA-PP1 to inhibit aPKC kinase activities, (Supplementary Figure 4I). PAR-3 transportation by cortical flow concentrates PAR-3 at the anterior cortex, so disruption of flow could perhaps account for the small changes in PAR-3 oligomerization observed in these experiments [68, 69]. Overall, the data suggest that aPKC kinase activity and aPKC kinase-substrate binding play, at most, a minor role in cooperative aPAR complex assembly.

### PAR-3 / aPKC cooperativity is developmentally regulated

PAR-3 oligomerization and cortical aPAR clustering are prevalent during the polarity establishment phase and decrease during maintenance phase (Figure 1C, Figure 6A) [45, 51, 64, 65, 68]. We, therefore, examined whether PAR-3 / aPKC cooperativity was altered during polarity maintenance using sc-SiMPull. During maintenance phase, the distribution of PAR-3 cluster sizes was no longer obviously biphasic, and it more closely matched a simple exponential distribution (Figure 7B), suggesting that PAR-3 / aPKC cooperativity was reduced. PAR-3 oligomers interacting with aPKC were significantly smaller in the maintenance phase than in the establishment phase (Figure 7C, E). We utilized cortical imaging with chimeric labeling to confirm these results in intact cells. Indeed, cortical PAR-3 oligomers were significantly smaller in the maintenance phase, consistent our sc-SiMPull data and with previous reports (Figure 7F-H) [45, 51, 64, 68]. Together, these data show that PAR-3 / aPKC cooperativity is developmentally regulated in the *C. elegans* zygote.

**Figure 7:**
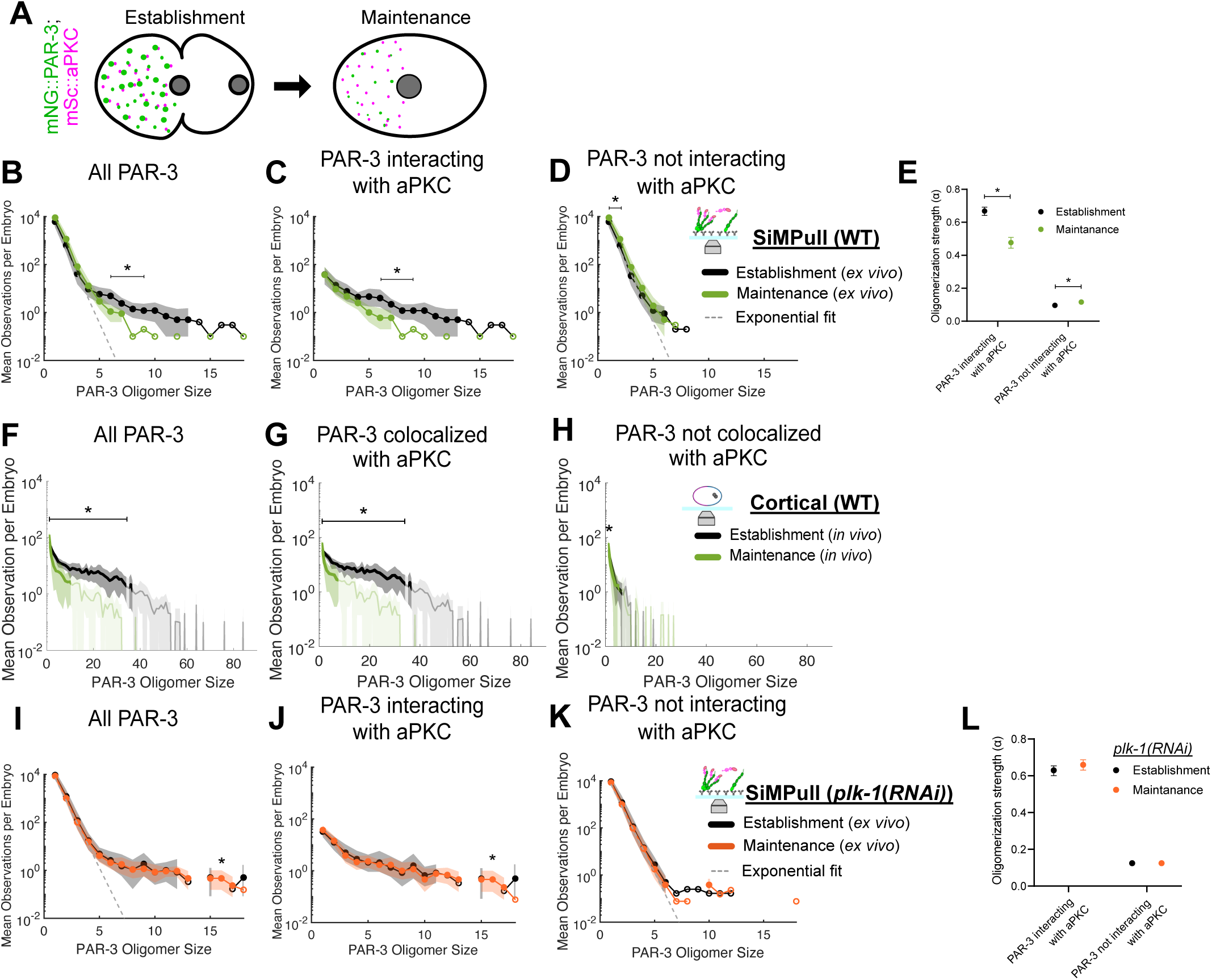
PLK-1 downregulates PAR-3 / aPKC cooperativity during the polarity maintenance phase. (A) Illustration of reduced cortical PAR-3 clustering during the maintenance phase. (B-E) Comparison of the oligomer size distributions for all PAR-3 (B), PAR-3 interacting with aPKC (C), and PAR-3 not interacting with aPKC (D) measured using sc-SiMPull. Labeling conventions for these plots are illustrated in Figure 2D. For clarity, only oligomer sizes up to 18 subunits are shown; the entire range of the data is plotted in Supplementary Figure 5A-C. * indicates significant differences between establishment- and maintenance- phase wild-type embryos (based on effect sizes plotted in Supplementary Figure 5D-F). The dashed line is an exponential fit from PAR-3 not interacting with aPKC in wild-type. The control dataset (black) is the same data as plotted in Fig. 5B-D and is shown again here for to facilitate comparison. (E) The oligomerization strength (α) from the exponential fit and its bootstrap 95% confidence interval. * indicates significant differences between groups because their confidence intervals do not overlap. n=67,479 PAR-3 complexes from N=10 embryos (Establishment); n=101,817 PAR-3 complexes from N=10 embryos (Maintenance). (F-H) Comparison of the oligomer size distributions for all PAR-3 (F), PAR-3 colocalized with aPKC (G), and PAR-3 not colocalized with aPKC (H) from cortical imaging with chimeric labeling in establishment or maintenance phase. The line graph shows the mean and its bootstrap 95% confidence interval. * indicates significant differences between establishment and maintenance (based on effect sizes plotted in Supplementary Figure 5G-I). The curve is truncated where it passes below 10^-2^; for these complex sizes, there were 0 observations in our dataset. The faint region indicates <20 total observations of a given size. The control dataset (black) is the same data as plotted in Fig. 5B-D and is shown again here for to facilitate comparison. n=3,099 PAR-3 clusters from N=10 embryos (Establishment); n=1,427 PAR-3 clusters from N=7 embryos (Maintenance). (I-L) Comparison of the oligomer size distributions for all PAR-3 (I), PAR-3 interacting with aPKC (J), and PAR-3 not interacting with aPKC (K) measured using sc-SiMPull. Labeling conventions for these plots are illustrated in Figure 2D For clarity, only oligomer sizes up to 18 subunits are shown; the entire range of the data is plotted in Supplementary Figure 5K–M. * indicates significant differences between establishment- and maintenance-phase *plk-1(RNAi)* embryos (based on effect sizes plotted in Supplementary Figure 5N-P). The dashed line is an exponential fit from PAR-3 not interacting with aPKC in wild-type. (L) The oligomerization strength (α) from the exponential fit and its bootstrap 95% confidence interval. * indicates significant differences between groups because their confidence intervals do not overlap. n=126,728 PAR-3 complexes from N=12 embryos (Establishment); n=123,768 PAR-3 complexes from N=13 embryos (Maintenance).

PLK-1 is a cell cycle regulator and is thought to dissociate PAR-3 oligomers by phosphorylating a site within the PAR-3 oligomerization domain during polarity maintenance phase [51]. To determine if PLK-1 regulates PAR-3 / aPKC cooperativity, we knocked down PLK-1 using RNAi and used sc-SiMPull to compare PAR-3 / aPKC complexes between establishment and maintenance-phase embryos. Strikingly, following PLK-1 depletion, both the PAR-3 / aPKC interaction and the distribution of oligomer sizes were identical between establishment and maintenance (Figure 7I-L). During maintenance phase, *plk-1(RNAi)* embryos had larger PAR-3 oligomers than wildtype, and these abnormally large oligomers contained aPKC (Supplementary Figure 6A-D). Thus, PLK-1 activity is necessary to downregulate PAR-3 / aPKC cooperativity during polarity maintenance. Overall, these data show that PAR-3 / aPKC cooperativity is developmentally regulated, and PLK-1 plays an essential role in dissociating PAR-3/ aPKC clusters during the maintenance phase.

## Discussion

The PAR-3 / aPKC / PAR-6 complex plays a critical role in cell polarity. Previous studies have shown that PAR-3 oligomerization and its interaction with aPKC are necessary to transport aPKC to one side of the cell during polarity establishment [51]. Our results showed that PAR-3 oligomerization and PAR-3 binding to aPKC / PAR-6 exhibit positive cooperativity. We identified genetic perturbations that specifically eliminate cooperativity without entirely blocking PAR-3 / aPKC / PAR-6 interactions. These mutations abolish polarity, demonstrating that cooperativity is required for polarity establishment. Cooperativity is a product of multivalency between PAR-3 and aPKC / PAR-6. We identified two key interactions: The PDZ / PDZ-ligand interaction between aPKC and PAR-3, and an interaction involving PAR-6. We propose that within an oligomeric PAR-3 / aPKC / PAR-6 complex, each aPKC / PAR-6 heterodimer makes physical contact with two different PAR-3 molecules, thereby achieving more stable binding while crosslinking and stabilizing the oligomer (Figure 8). Whether aPKC’s kinase domain interacts stably with its substrate on PAR-3 *in vivo* has been controversial [39, 44, 47], but we present evidence that this interaction plays at most a minor role in PAR-3 / aPKC / PAR-6 cooperativity. Lastly, PAR-3 / aPKC / PAR-6 cooperativity is downregulated during polarity maintenance, and PLK-1 regulates this downregulation. Based on these findings, we conclude that multivalent interactions between PAR-3 and aPKC / PAR-6 cooperatively assemble the aPAR complex to enable polarity establishment. In polarity maintenance, PLK-1 activation quenches PAR-3 / aPKC cooperativity (Figure 8) to prevent aberrant PAR-3 segregation due to cortical flows that occur during cytokinesis [51].

**Figure 8:**
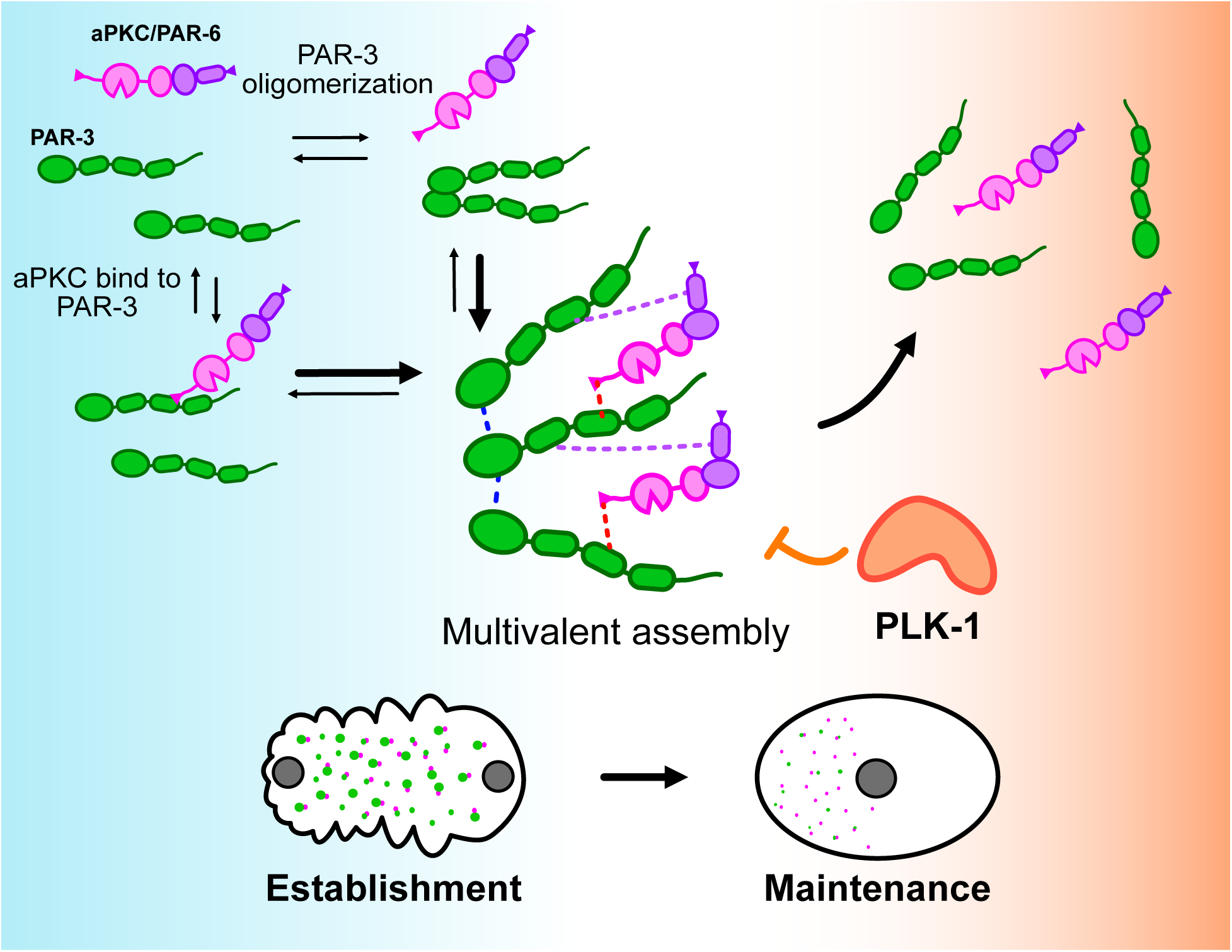
Conclusion. The illustration of the main concept of this study: (1) PAR-3 oligomerization cooperates with PAR-3 / aPKC interaction to form PAR complexes. (2) The PAR-3 / aPKC cooperativity depends on multivalency, with both aPKC and PAR-6 making interactions with PAR-3. (3) The cooperativity is prevalent in establishment embryos, but PLK-1 downregulates the cooperativity during maintenance.

The multivalent assembly process we propose for PAR-3 / aPKC / PAR-6 clusters is somewhat reminiscent of biomolecular condensation, wherein multivalent, often disordered proteins undergo phase separation into liquid-or gel-like condensates [77]. However, PAR-3 / aPKC / PAR-6 clusters are different from biomolecular condensates in several important respects. First, our analysis of their size distribution (Figure 2F) suggests that aPAR clusters are linear oligomers rather than disordered condensates, in agreement with structural data [50]. We observed monotonically decreasing, exponential size distributions, which matches the prediction for linear oligomers [68, 78] but is inconsistent with the log-normal size distributions observed for phase-separated protein droplets [79]. Second, the interactions involved in aPAR cluster assembly are relatively high-affinity interactions involving well-structured domains [37, 39, 47, 80–82], in contrast to the low-affinity interactions between disordered regions that are commonly associated with condensates. Third, aPAR clusters are not liquid-like, because they persist for several minutes following extraction from cells. We therefore think of aPAR clusters as well-ordered, stable assemblies rather than condensates. In this regard, it is interesting that a fragment of mammalian Par3 has been reported to undergo phase separation *in vitro,* both alone or in combination with Par6β [83]. That study involved unrealistically high concentrations of PAR protein fragments *in vitro,* and overexpression in mammalian cells; it did not examine the intact aPAR complex at endogenous expression levels, as we have done. Therefore, although we do not exclude the possibility that PAR proteins may phaseseparate at higher concentrations, we do not think that such behavior is relevant to their normal physiological function.

PAR-3 / aPKC cooperativity modulates the lipid binding affinity of the aPAR complex. We found that cortical PAR-3 is mostly colocalized with aPKC, suggesting that most monomeric and dimeric PAR-3 resides in the cytosol (Figure 3C). This is consistent with our previous work showing that 3mers of PAR-3 were necessary and sufficient for membrane binding and normal polarity establishment [54]. The size distributions of PAR-3 clusters interacting with aPKC are similar between *ex vivo* and *in vivo* assays, but smaller PAR-3 clusters not interacting with aPKC were rarely observed on the membrane *in vivo* (Figure 2F, 3C). In addition, the aPKC(V597A) mutant decreases cortical localization of PAR-3 and aPKC (Figure 4F, Supplementary Figure 2F), suggesting that PAR-3 / aPKC cooperativity is crucial for aPAR complex binding to the membrane. Previous studies found that PAR-3 binds to the membrane via its PDZ domain [40] and coiled-coil C-terminus [41–43], and aPKC also contains a membrane associate domain [84]. Thus, assembly of PAR-3 / aPKC clusters is expected to result in much stronger membrane binding due to avidity caused by multiple membrane binding sites within the complex.

PAR-6 can support PAR-3 / aPKC cooperativity by binding to both aPKC and PAR-3 In our sc-SiMPull data, bigger PAR-3 oligomers have a higher chance of containing aPKC and PAR-6 (Figure 2C). PAR-6 knockdown downregulates PAR-3 / aPKC cooperativity (Figure 5), confirming that aPKC and PAR-6 are indispensable partners in aPAR complex formation [34, 57]. The exact nature of PAR-6 interactions within this complex remain somewhat unclear. Several studies suggest PAR-3 / PAR-6 interaction via the PAR-6 PDZ domain or its PDZ ligand binding to the PAR-3 PDZ domain [34, 37, 57]. However, the PAR-6 PDZ ligand is not conserved across species [39], and a PAR-6 PDZ ligand mutant has a weaker defect in *Drosophila* neuroblast polarity compared to an aPKC PDZ-ligand mutant [56], implying that the PAR-3 / PAR-6 interaction is redundant or otherwise dispensable in that system. However, even a weak interaction could contribute to multivalency for PAR-3 /aPKC / PAR-6 cooperativity. It also remains possible that PAR-6 interacts with PAR-3 in a way that does not require a PDZ / PDZ ligand interaction. A third hypothesis is that PAR-6 activates cooperativity by binding to aPKC inducing a conformational change that allows the aPKC kinase domain to make a second interaction with PAR-3 [84–86]. To sum up, PAR-6 regulates multivalent PAR-3 / aPKC assembly, but we still do not know whether this occurs via direct PAR-3 / PAR-6 contacts or aPKC activation.

PAR-3 contains an aPKC substrate peptide (sometimes called CR3), and previous studies have shown that PAR-3 with CR3 mutants have reduced PAR-3 / aPKC interactions in mammalian systems [31, 32, 61]. However, both phospho-mimetic or phospho-defective mutants at CR3 of PAR-3 can substitute wild-type PAR-3 functions in polarity establishment in *C. elegans* zygotes, implying that either aPKC does not phosphorylate PAR-3 CR3 in *C. elegans*, or this phosphorylation is dispensable for PAR-3 function in the zygote [52]. We found no clear evidence that aPKC / PAR-3 CR3 interactions contribute to aPAR complex assembly, either *ex vivo* or *in vivo* (Figure 6). A caveat is that aPKC not only recognizes the CR3 but also nearby FR and KR domains to enhance substrate-kinase binding [44], and therefore, ATP may not completely dissolve the interaction between aPKC and full-length PAR-3 resulting from substrate-kinase binding in our sc-SiMPull assays. Nevertheless, we would have expected to see some effect of ATP addition if kinase-substrate binding were critical for cooperativity. On the other hand, cortical imaging with chimeric labeling showed small differences in the abundance of PAR-3 clusters on the membrane, especially an increase in the abundance of cortical 2mers and 3mers, upon 1NA-PP1 treatment (Figure 6E, Supplementary Figure 4H). These changes may be due to disruption of cortical flow or another indirect consequence of aPKC inhibition.

Our previous work has shown that PLK-1 downregulates PAR-3 oligomerization during polarity maintenance phase. PLK-1 could directly phosphorylate a fragment of PAR-3 *in vitro,* and a phosphomimetic mutant in PAR-3 eliminated PAR-3 oligomerization [51]. Therefore, it seemed likely that in cells, PLK-1 down-regulated PAR-3 clusters by directly phosphorylating PAR-3 and reducing the concentration of oligomerization-competent PAR-3 monomers. However, the results we present here argue against this simple model. If PLK-1 acted by reducing the amount of oligomerization-competent PAR-3 during maintenance phase, then the abundance of oligomers containing and lacking aPKC should both have been affected. In contrast to this prediction, we found that oligomerization of PAR-3 / aPKC complexes was strongly reduced during maintenance phase, but oligomerization of PAR-3 alone (lacking aPKC) was essentially unchanged (Figure 7C-D). Furthermore, loss of PLK-1 did not cause large changes in oligomerization of PAR-3 clusters lacking aPKC at any stage of the cell cycle (Supplementary Figures S6C and S6I). Together, these observations suggest that PLK-1 phosphorylation of PAR-3 does not simply quench PAR-3 oligomerization by reducing the available monomer pool. Nor does PLK-1 directly affect the PAR-3 / aPKC interaction, because interaction of PAR-3 monomers and dimers with aPKC is unaltered during maintenance phase and unaffected by PLK-1 depletion (Figures 7C, 7I, S6B and S6H). Instead, PLK-1 appears to specifically regulate PAR-3 / aPKC cooperativity. How this occurs mechanistically remains unclear, but one possibility is that PLK-1 is specifically recruited to aPAR clusters and contributes to their active disassembly at the establishment-maintenance transition. Future studies will address this and other hypotheses for PLK-1’s mechanism of action.

In yeast polarity, the cell cycle regulator CDK couples with CDC-42, a conserved Rho-family GTPase, to coordinate cell polarity and cell cycle [87]. CDC-42 also regulates anterior polarity in *C. elegans* during polarity maintenance [45, 88–90]. Interestingly, CDC-42 competes with PAR-3 for aPKC / PAR-6 binding *in vitro* [86, 91], and we previously showed that CDC-42 / PAR-3 / aPKC / PAR-6 tetramers are not detectably present in the zygote. CDC-42 gains stronger binding affinity to PAR-6 during polarity maintenance, suggesting that CDC-42 may contribute to displacing PAR-6 / aPKC from PAR-3 and reducing cooperative PAR complex assembly during polarity maintenance.

Embryonic development relies on dynamic regulation of protein-protein interactions to control cell behavior. Dissecting key signaling interactions biochemically is therefore critical to understand how organisms are constructed. However, the rapid growth and development of early embryos makes biochemistry difficult because bulk sample collection is often impossible, and heterogeneity can obscure rapid regulation at specific developmental stages. Taking *C. elegans* zygotes as an example, the time from symmetry breaking to the first mitosis is only 15 minutes. Previous studies have shown that the biochemistry of PAR complexes significantly changes within this short period of time [51, 92], emphasizing that studying mixed-stage samples, even if they are pure populations of zygotes, cannot reveal developmentally important changes in the interactions of key signaling molecules. In this study and others from our group [58, 92], we performed single-cell, single-molecule biochemistry to overcome this obstacle and investigate how protein-protein interactions are remodeled during normal embryonic development. sc-SiMPull allows us to examine the biochemistry of full-length PAR-3 and to resolve changes in its interactions that occur within minutes *in vivo.* We confirmed the results using *in vivo* single-molecule cortical imaging with chimeric labeling. Combining these *in vivo* and *ex vivo* assays, along with mutant, knockdown, and drug conditions, allowed us to elucidate how PAR-3 recruits aPKC / PAR-6 to form anterior complexes during polarity establishment. This work highlights the potential of single-molecule biochemistry as applied in developmental biology.

### Limitations of the Study

First, we limited our investigation to the *C. elegans* zygote, so we do not exclude the possibility that aPAR complex assembly may occur differently or be regulated in another way in other developmental stages or other systems. Second, due to embryonic lethality, we were unable to isolate truncation mutants of PAR-6 that would have narrowed down the regions of the protein involved in cooperativity; we were therefore limited to RNAi approaches that deplete the total protein levels. *In vitro* approaches might be necessary to further pinpoint the binding sites within PAR-6 that contact PAR-3. Third, our experiments to address the effects of aPKC kinase activity on aPAR complex cooperativity yielded mainly negative results, so we cannot formally exclude the possibility that stronger perturbations or sensitized backgrounds would have revealed a clear effect. Finally, a fundamental limitation of all biochemical approaches is that they require removal of biological macromolecules from the native cellular environment. sc-SiMPull mitigates this limitation by counting protein complexes within seconds after cell lysis, but no biochemical approach can avoid the caveat that protein interactions may be disrupted or altered by cell lysis.

## Author Contributions

D.J.D. conceived of the project, supervised the work and secured funding. S-P.H. and D.J.D. performed the experiments, analyzed the data, and co-wrote the manuscript.

## Acknowledgments

We thank Luke Lavis for sharing JaneliaFluor dyes, Nathan Goehring for sharing *C. elegans* strains, and Edwin Munro and members of the Dickinson laboratory for helpful discussions and comments on the manuscript. This work was supported by a fellowship from the Ministry of Education, Republic of China (Taiwan) (S-P.H.); and by NIH R01 GM138443, NIH R21 GM144817 and NSF MCB 2237451 (D.J.D.).

---

**Supplementary Figure 1, related to Figures 2-3:**
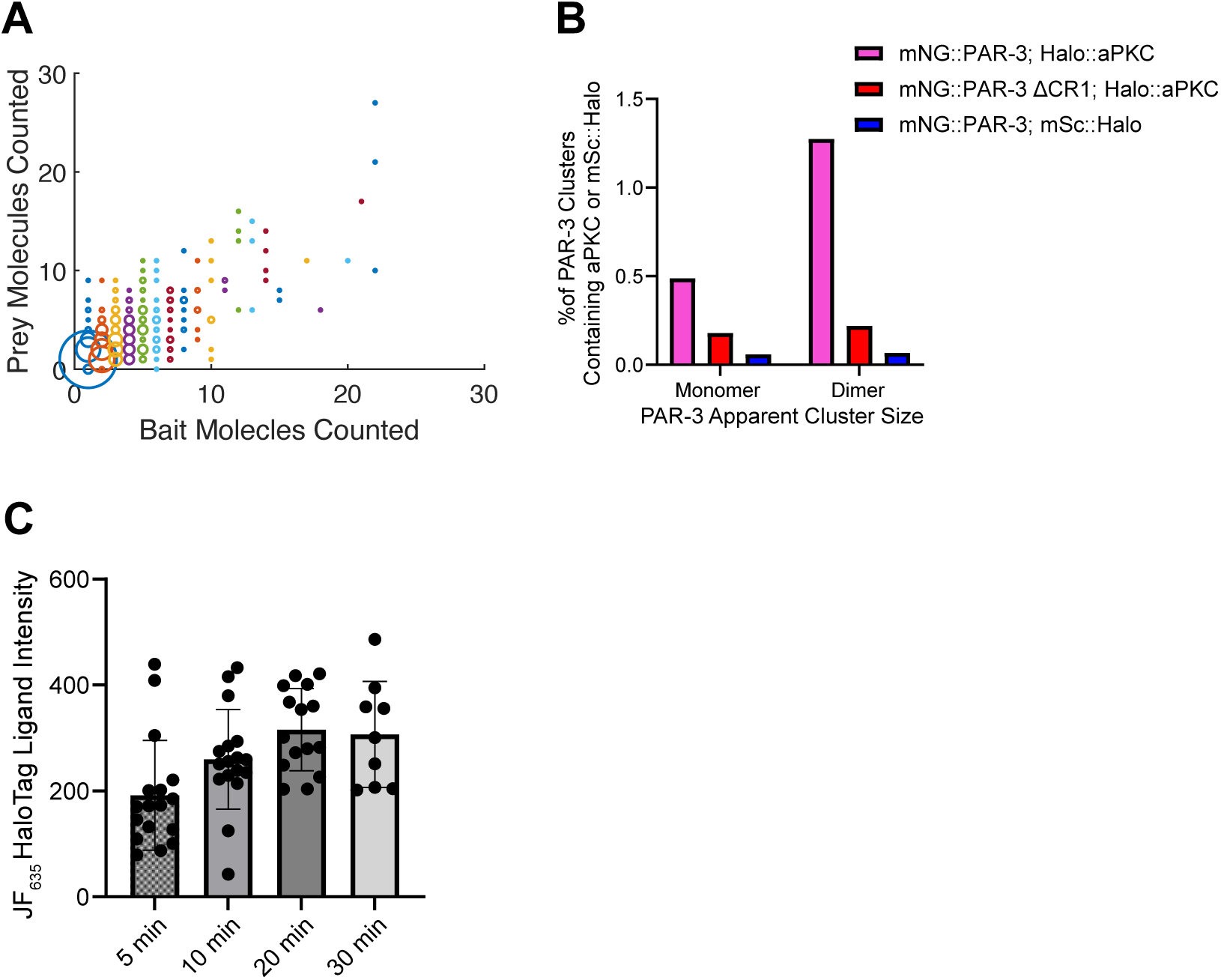
Additional data regarding aPAR cluster assembly during establishment phase. (A) Comparison of numbers of Halo::aPKC and mNG::PAR-3 molecules per complex, based on two-color photobleaching step counting in sc-SiMPull experiments. The size of each circle indicates the number of observations found in each position. n=158,514 PAR-3 complexes from N=26 embryos. (B) Fraction of PAR-3 monomers and dimers in complex with Halo::aPKC or mSc::Halo (negative control) measured using sc-SiMPull. n=37,747 PAR-3 monomers and 2,430 PAR-3 dimers from N=13 embryos (mNG::PAR-3; Halo::aPKC); n=54,217 PAR-3 monomers and 3,189 dimers from N=13 embryos (mNG::PAR-3 ΔCR1; Halo::aPKC); n=17,046 monomers and 1,493 dimers from N=8 embryos (mNG::PAR-3; mSc::Halo). (C) Time course of Halo dye staining of permeabilized embryos for cortical imaging with chimeric labeling. The replicates are: 9 embryos for 5-min staining, 9 embryos for 10-min staining, 8 embryos of 20-min staining, and 5 embryos of 30-min staining

**Supplementary Figure 2, related to Figure 4:**
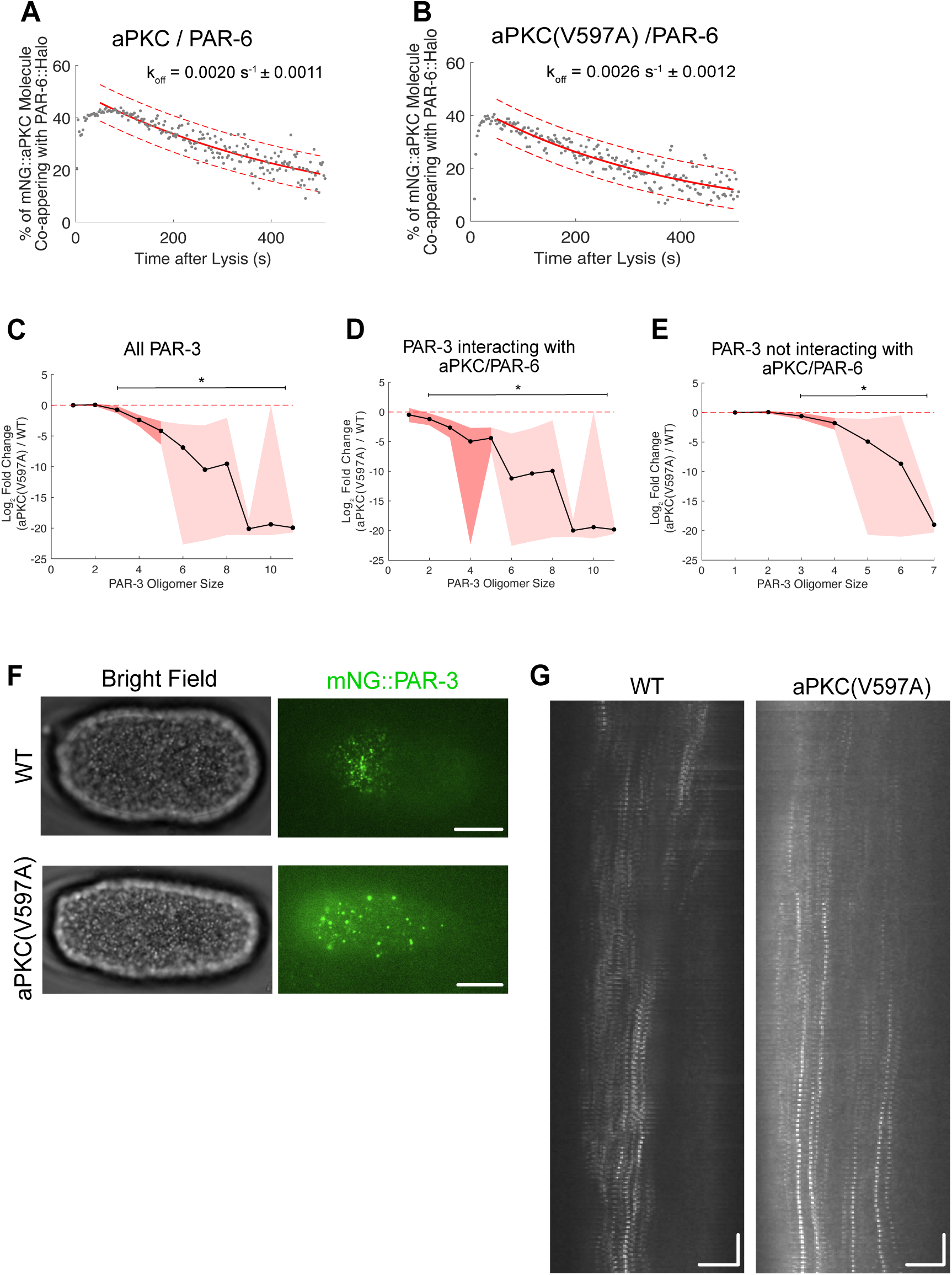
Controls and cortical flow phenotype for aPKC(V597A) mutant. (A-B) Fraction of mNG::aPKC molecules co-appearing with PAR-6::Halo as a function of time since cell lysis. The red curves are fits to single-exponential decay functions, used to estimate koff, [58] and the dashed lines show the 95% confidence interval. n=158,802 aPKC / PAR-6 complexes from N=11 embryos (WT); n=108,675 aPKC / PAR-6 complexes from N=12 embryos (aPKC (V597A)). (C-E) Effect size plots for the sc-SiMPull experiments in Figure 4B-D. The comparison is over all PAR-3 (A), PAR-3 interacting with aPKC / PAR-6 (B), and PAR-3 not interacting with aPKC / PAR-6 (C). The line shows the mean, and the colored area shows a bootstrap 95% confidence interval. The line plot only shows sizes with enough observations to calculate a confidence interval, and the faint shaded region indicated only one dataset had enough observations (as defined in Methods). * indicates the region considered significant, which we define by two criteria: (1) The 95% confidence interval of the effect size does not overlap with 0, and (2) at least one genotype had a sufficient number of observations of a given oligomer size to allow calculation of a bootstrap 95% confidence interval (see Methods). (F) Cortical images of mNG::PAR-3 in wild-type and aPKC(V597A) embryos at pronuclear meeting. Anterior is to the left. Scale bars represent 10 µm. (G) Kymograph of cortical mNG::PAR-3 in wild-type and aPKC(V597A) embryos. Anterior is to the left. The horizontal scale bars represent 10 µm. The vertical scale bars represent 30 seconds.

**Supplementary Figure 3, related to Figure 5:**
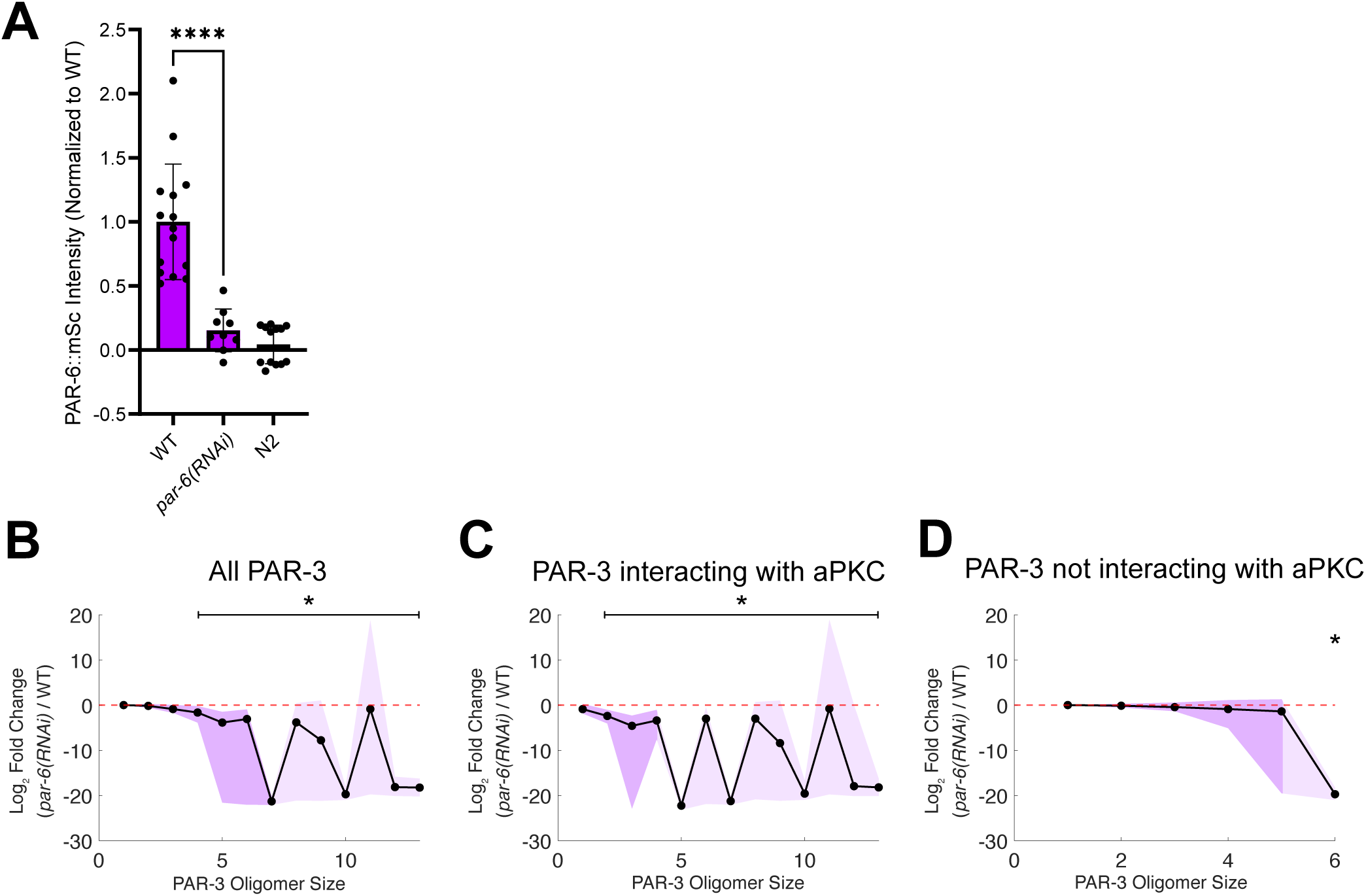
PAR-6 depletion affects cooperative aPAR complex assembly. (A) PAR-6::mSc intensity in an untreated knock-in strain, in the knock-in strain after *par-6* RNAi, and in the non-tagged strain N2 (used as a control for autofluorescence). The intensity is normalized to the average without RNAi. **** indicates a significant difference between WT and *par-6(RNAi)* by Student’s t test. N=14 embryos (WT); N=9 embryos (*par-6(RNAi)*); N=13 embryos (N2). (B-D) Effect size plots for the sc-SiMPull experiments in Figure 5B-D. The comparison is over all PAR-3 (B), PAR-3 interacting with aPKC (C), and PAR-3 not interacting with aPKC (D). The line shows the mean, and the colored area shows a 95% confidence interval after bootstrapping. The line plot only shows sizes with enough observations to calculate a confidence interval, and the faint shaded region indicated only one dataset had enough observations (as defined in Methods). * indicates the region considered significant, which we define by two criteria: (1) The 95% confidence interval of the effect size does not overlap with 0, and (2) at least one genotype had a sufficient number of observations of a given oligomer size to allow calculation of a bootstrap 95% confidence interval (see Methods).

**Supplementary Figure 4, related to Figure 6:**
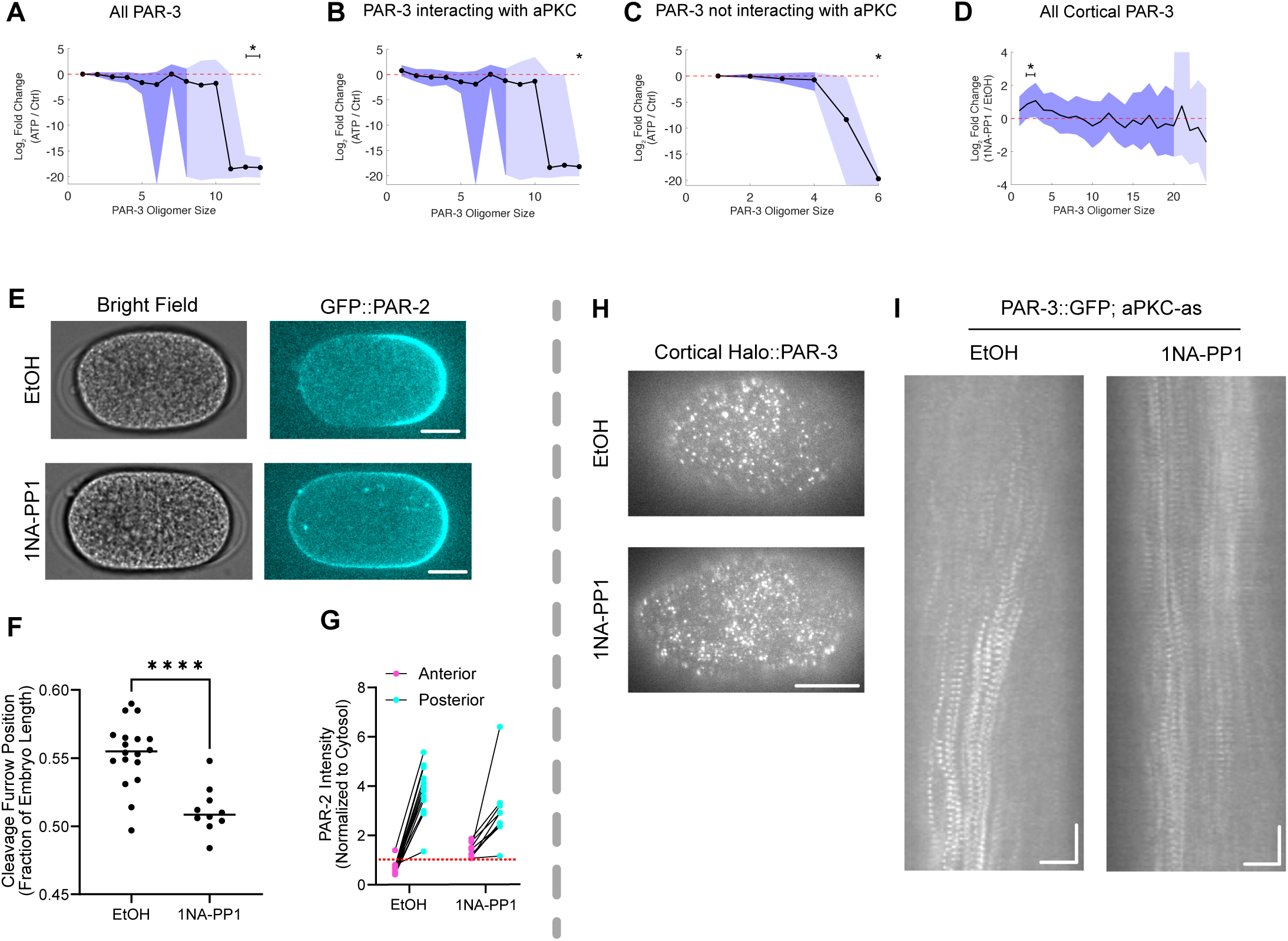
aPKC kinase activity has at most a minor effect on aPAR complex assembly. (A-C) Effect size plots for the sc-SiMPull experiments in Figure 6B-D. The comparison is over all PAR-3 (A), PAR-3 interacting with aPKC (B), and PAR-3 not interacting with aPKC (C). The line graph shows the mean, and the colored area shows a 95% confidence interval after bootstrapping. The line plot only shows sizes with enough observations to calculate a confidence interval, and the faint shaded region indicated only one dataset had enough observations (as defined in Methods). * indicates the region considered signficiant, which we define by two criteria: (1) The 95% confidence interval of the effect size does not overlap with 0, and (2) at least one condition had a sufficient number of observations of a given oligomer size to allow calculation of a bootstrap 95% confidence interval (see Methods). (D) Effect size plots for the cortical imaging experiment with chimeric labeling, shown in Figure 5F. The line shows the mean, and the colored area shows a bootstrap 95% confidence interval. The line plot only shows sizes with enough observations to calculate a confidence interval, and the faint shaded region indicated only one dataset had enough observations (as defined in Methods). * indicates the region considered significant, as in (A-C). The replicates are: N=18 embryos (EtOH); N=10 embryos (1NA-PP1). (E) Images PAR-2::GFP localization in aPKC-as embryos following treatment with ethanol or 1NA-PP1. Anterior is to the left. Scale bars represent 10 µm (F) Length of the AB cell as a fraction of total embryo length in 2-cell aPKC-as embryos treated with ethanol control or 1NA-PP1. **** indicates a significant difference (Student’s t-test). N=13 embryos (EtOH); N=10 embryos (1NA-PP1). (G) Quantification of anterior and posterior PAR-2::GFP intensity following ethanol or 1NA-PP1 treatment. The intensity is normalized to cytosol intensity. The red dashed line indicates the value of 1, which indicates no measurable cortical enrichment. (H) Cortical images of Halo::PAR-3 in aPKC-as embryos following treatment with ethanol or 1NA-PP1. Anterior is to the left. Scale bars represent 10 µm. (I) Kymographs of cortical PAR-3::GFP. The length of the images is around 7 minutes. Anterior is to the left. The horizontal scale bars represent 10 µm. The vertical scale bars represent 30 seconds.

**Supplementary Figure 5, related to Figure 7:**
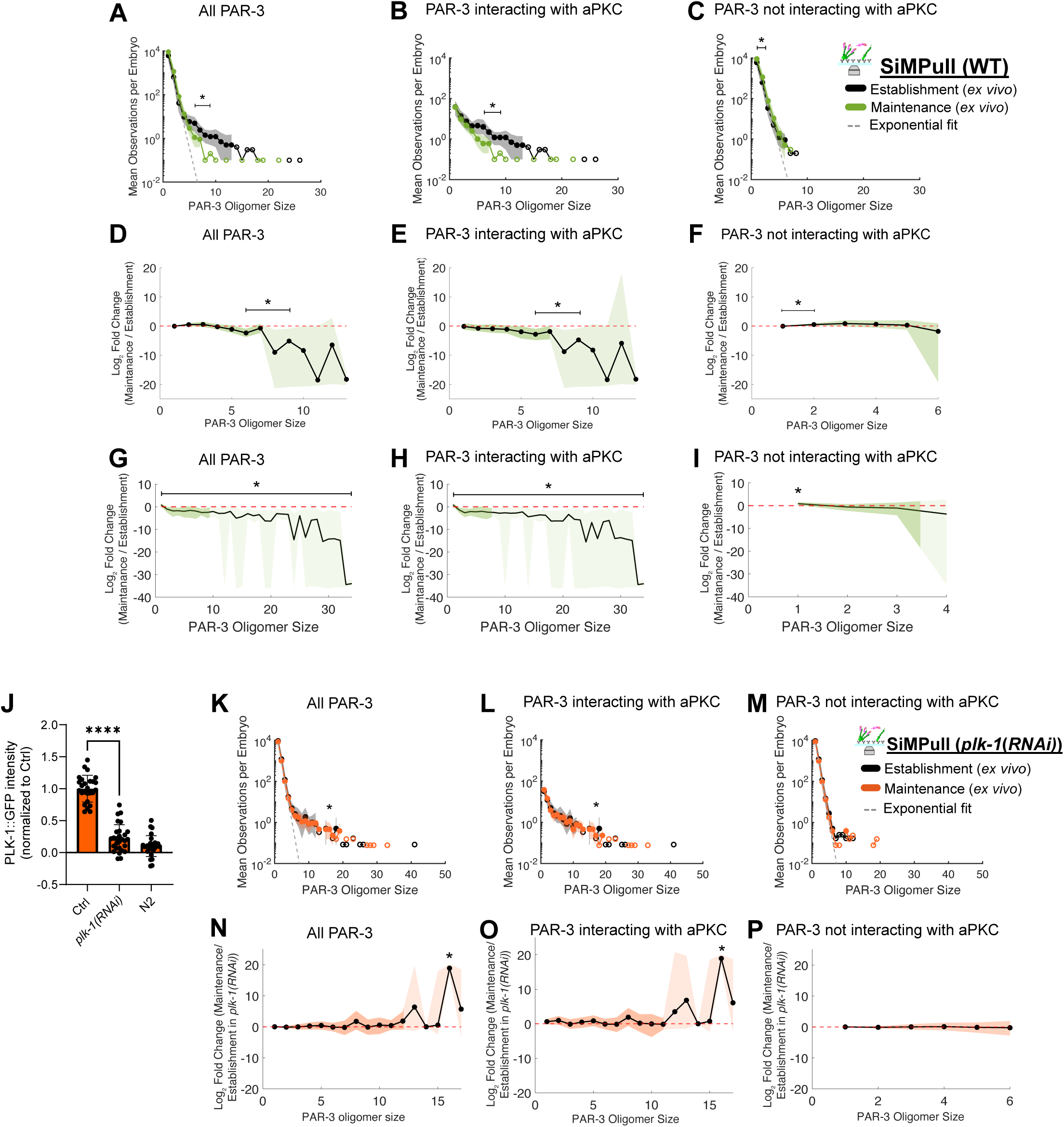
aPAR complex assembly in Establishment phase compared to Maintenance phase. (A-C) Full-scale plots of the data shown in Figure 7B-D (D-F) Effect size plots for the sc-SiMPull experiments in Figure 7B-D. The comparison is over all PAR-3 (D), PAR-3 interacting with aPKC (E), and PAR-3 not interacting with aPKC (F). The line graph shows the mean, and the colored area shows a bootstrap 95% confidence interval. * indicates the region considered significant, which we define by two criteria: (1) The 95% confidence interval of the effect size does not overlap with 0, and (2) at least one stage had a sufficient number of observations of a given oligomer size to allow calculation of a bootstrap 95% confidence interval (see Methods). (G-I) Effect size plots for the chimeric labeling experiments in Figure 7F-H. The comparison is over all PAR-3 (G), PAR-3 colocalized with aPKC (H), and PAR-3 not colocalized with aPKC (I). The line graph shows the mean, and the colored area shows a 95% confidence interval after bootstrapping. * indicates the region considered significant, as in A-C. (J) PLK-1::GFP intensity in an untreated knock-in strain, in the knock-in strain after *plk-1* RNAi, and in the non-tagged strain N2 (used as a control for autofluorescence).The intensity is normalized to the average without RNAi. **** indicates a significant difference between WT and *plk-1(RNAi)* by Student’s t-test. The data are from 26 replicates of WT, 28 replicates of *plk-1(RNAi)*, and 28 replicates of N2. (K-M) Full-scale plots of the data shown in Figure 7I-K (N-P) Effect size plots for the sc-SiMPull experiments in Figure 7I-K. The comparison is over all PAR-3 (N), PAR-3 interacting with aPKC (O), and PAR-3 not interacting with aPKC (P). The line graph shows the mean, and the colored area shows a 95% confidence interval after bootstrapping. * indicates the region considered significant, as in D-F.

**Supplementary Figure 6, related to Figure 7:**
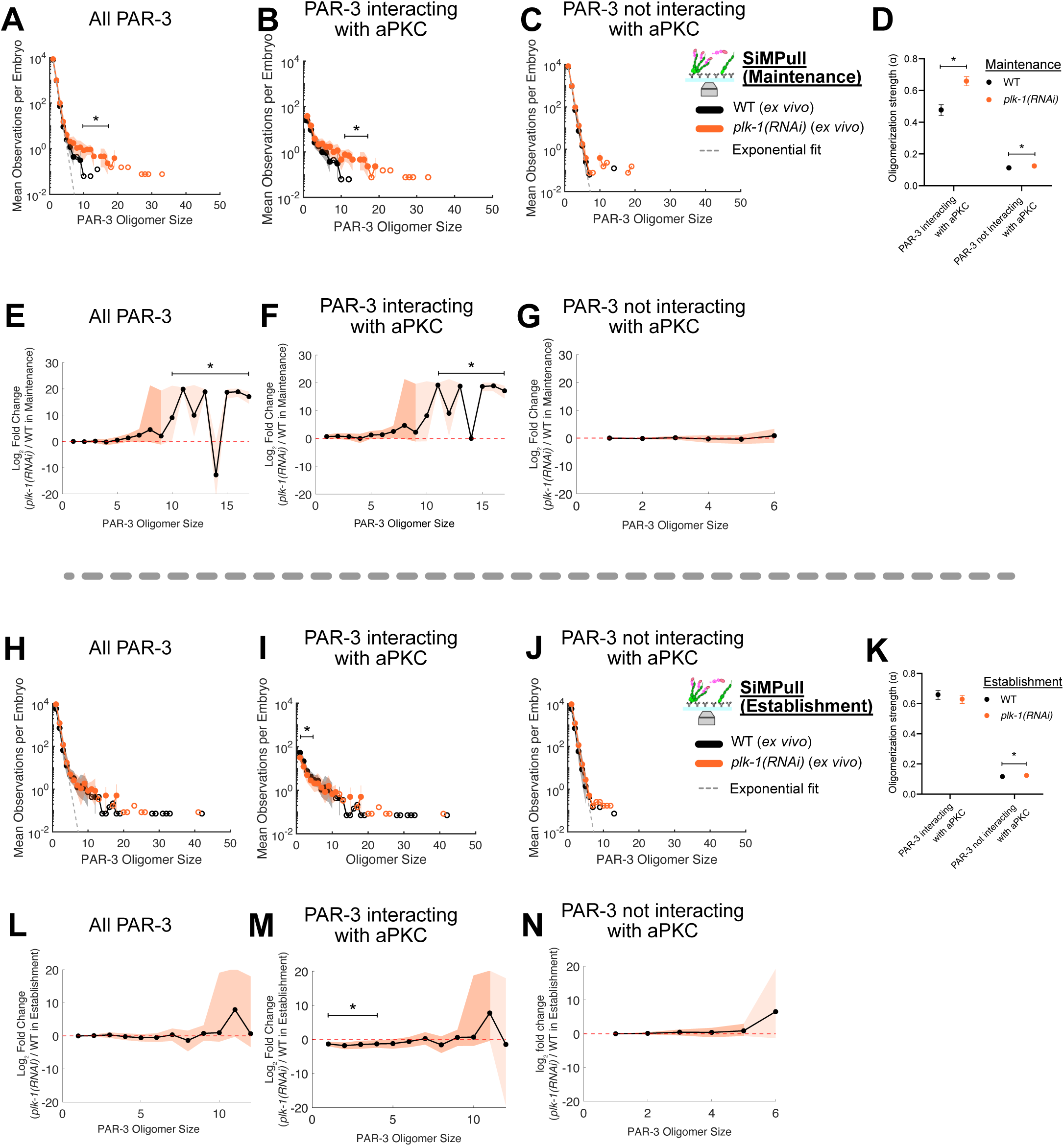
aPAR complex assembly in wild-type compared to *plk-1(RNAi)* (A-D) Comparison of the oligomer size distributions for all PAR-3 (A), PAR-3 interacting with aPKC (B), and PAR-3 not interacting with aPKC (C) measured using sc-SiMPull from Maintenance-phase embryos. Labeling conventions for these plots are illustrated in Figure 2D. * indicates significant differences between wild-type and *plk-1(RNAi)* embryos (based on effect sizes plotted in (E-G)). The dashed line is an exponential fit from PAR-3 not interacting with aPKC in wild-type establishment. (D) The oligomerization strength (α) from the exponential fit and its bootstrap 95% confidence interval. * indicates significant differences between groups because their confidence intervals do not overlap with each other after bootstrapping iterations. n=141,272 PAR-3 complexes from N=16 embryos (WT); n=123,768 PAR-3 complexes from N=13 embryos (*plk-1(RNAi)*). (E-G) Effect size plots for the sc-SiMPull experiments in (A-C). The comparison is over all PAR-3 (E), PAR-3 interacting with aPKC (F), and PAR-3 not interacting with aPKC (G). The line graph shows the mean, and the colored area shows its bootstrap 95% confidence interval. * indicates the region considered significant, which we define by two criteria: (1) The 95% confidence interval of the effect size does not overlap with 0, and (2) at least one genotype had a sufficient number of observations of a given oligomer size to allow calculation of a bootstrap 95% confidence interval (see Methods). (H-K) Comparison of the oligomer size distributions for all PAR-3 (H), PAR-3 interacting with aPKC (I), and PAR-3 not interacting with aPKC (J) measured using sc-SiMPull from Establishment-phase embryos. Labeling conventions for these plots are illustrated in Figure 2D. * indicates significant differences between wild-type and *plk-1(RNAi)* embryos (based on effect sizes plotted in (L-N)). The dashed line is an exponential fit from PAR-3 not interacting with aPKC in wild-type establishment. (K) The oligomerization strength (α) from the exponential fit and its bootstrap 95% confidence interval. * indicates significant differences between groups because their confidence intervals do not overlap. n=91,287 PAR-3 complexes from N=14 embryos (WT); n=126,728 from N=12 embryos (*plk-1(RNAi)*). (L-N) Effect size plots for the sc-SiMPull experiments in (H-J). The comparison is over all PAR-3 (L), PAR-3 interacting with aPKC (M), and PAR-3 not interacting with aPKC (N). The line graph shows the mean, and the colored area shows a 95% confidence interval after bootstrapping. * indicates the region considered significant, as in (E-G).

## Materials and methods

### C. elegans strains, material, organism, and software

*C. elegans* strains were cultured on NGM plates with *E. coli* OP50 for feeding, according to standard protocols. *C. elegans* strains containing fluorescent tags or mutations were constructed using established protocols for CRISPR/Cas9 gene editing [93]. All strains and materials are listed in supplementary table 1.

### Fabrication of microfluidic with antibodies

Homemade microfluidic devices for sc-SiMPull were fabricated exactly as described in a published protocol [94]. Briefly, we first vigorously mixed polydimethylsiloxane (PDMS, VWR, MSPP-DC2065622) and curing reagent at a 10:1 ratio, followed by pouring the PDMS mix onto a homemade silicon mold. PDMS was degassed by vacuum, spin-coated at 300 rpm, and baked at 85°C for 30 mins. Before device assembly, we treated the PDMS with air plasma for 30 sec and coverslips (VWR, 63769-01) in a UV-Ozone cleaner for 20 mins to form a permanent bond between PDMS and coverslip. Afterward, we treated the device with mPEG-Silane (Gelest, SIM6492.72-25GM) containing 0.02% biotin-PEG and 2% HPLC water for 30 mins, followed by washout 3 times with HPLC water. Devices were put in the box with desiccant at room temperature at least overnight (and up to 2 months) to cure the PEG layer before use. During the day of sc-SiMPull, we first rinsed the device with SiMPull buffer (10 mM Tris pH 8.0, 150 mM NaCl, 0.5% TX-100, and 0.1 mg/mL BSA). Afterward, we coated the channel with 0.2 mg/mL neutravidin (ThermoFisher Scientific, 31000) for 10 mins, followed by biotinylated mNG nanobodies for 10 mins. After washout extra neutravidin and nanobodies, we further conducted sc-SiMPull

### Labeling Halo tags with ligand dye

We used liquid culture to label the HaloTag for sc-SiMPull. Liquid cultures of *E. coli* HB101 or OP50 were spun down and resuspended in 100 µL S media, containing 100 mM NaCl, 6 mM K_2_HPO_4_, 44 mM KH_2_PO_4_, 13 µM cholesterol, 10 mM potassium citrate (pH 6.0), 3 mM CaCl_2_, 3 mM MgCl_2_, 65 µM EDTA, 25 µM FeSO_4_, 10 µM MnCl_2_, 10 µM ZnSO4, 1 µM CuSO4. JF_646_ HaloTag ligand [95], dissolved in DMSO, was added to the bacterial mix to a final concentration of 1.5 µM. We picked 30-60 L4 worms into 30 µL of the dye/food mix and incubated them at 20°C on a 250-rpm rotor for 16-24 hr.

For cortical imaging with chimeric labeling, we first treated the L4 worms with *perm-1* RNAi bacteria [96] for 16-24 hr. We mixed JF_635_ and JF_585_ HaloTag ligands [97] at a 100:1 ratio. Afterward, we dissected the *perm-1* depleted worms in Shelton’s growth media (0.5 mg/L inulin, 20 mg PVP powder, 1% BWE vitamins, 1% lipid concentrate, 1X pen/strap in Schneider’s drosophila media with 35% FBS) containing a total concentration of 1.5 µM HaloTag ligand dye, followed by incubation 10-20 mins (Supplementary Figure 1C). Afterward, we washed the embryo with SGM (without dye) twice before imaging.

### RNA interference

RNA interference (RNAi) was carried out by feeding using *E. coli* clones from an RNAi feeding library [98]. To knock down *perm-1* for drug treatment and HaloTag labeling in cortical images with chimeric labeling, we first concentrated a 5 mL *E. coli* overnight culture producing *perm-1* RNAi into 1 mL and add 50-100 µL bacteria to an LB plate with 1mM IPTG and 25 µg/mL carbenicillin, followed by leaving the plate overnight to express dsRNA. We picked 30-60 L4 worms onto the RNAi plate and incubated them at 20°C for 16-24 hr before use. To knock down *par-6* and *plk-1* for sc-SiMPull, we first cultured *E. coli* producing *par-6* or *plk-1* RNAi in liquid LB overnight to reach stationary phase and treated with 5 mM IPTG for 4 hr. We put L1 worms on an IPTG plate with RNAi bacteria to increase knockdown efficiency. Afterward, we incubated the 30-60 L4 worms in S medium containing dsRNA-producing *E. coli*, 1.5 µM HaloTag ligand, 1 mM IPTG, and 25 µM carbenicillin at 20°C for 16-24 hr.

### Single-cell single-molecule pulldown (sc-SiMPull)

sc-SiMPull was carried out as previously described [58]. Briefly, we first washed an antibody-functionalized microfluidic channel once with SiMPull buffer. Worms were dissected in egg buffer (118 mM NaCl, 40 mM KCl, 3.4 mM MgCl_2_, 3.4 mM CaCl_2_, and 5 mM HEPES pH 7.4). An embryo of the appropriate stage was selected, washed twice with SiMPull buffer, and further transferred to the inlet of the SiMPull device. We used a 27G needle to push the embryo into the middle of the channel and then sealed the inlet and outlet with clear tape. The embryo was transferred to the TIRF microscope and lysed using a shot from a 1064 nm pulsed laser. Immediately after lysis, we recorded multicolor single-molecule images 50-60 µm away from the lysis point. We acquired the single-molecule images using a home-built total internal reflection fluorescence (TIRF) microscope. The microscope was constructed around a MadCity Labs RM21 advanced platform and equipped with 488 nm, 561 nm and 638 nm lasers for fluorescence excitation; a 60x, 1.49 NA TIRF objective lens (Olympus); micromirrors for TIRF excitation [99]; a home-built 4-color image splitter for simultaneous multicolor acquisition [100]; a Photometrics PrimeBSI Express camera; and controlled using MicroManager software with custom acquisition plug-ins. The exposure time was 50ms and 10000 frames were captured in each experiment. The following laser powers were used: 10-20 mW 488 nm laser for mNG; 40 mW 561 nm laser for mScarlet-I; and 50-60 mW 638 nm laser for JF_646_. In the ATP addition experiment, we used SiMPull buffer with the addition of 1 mM ATP and 5 mM MgCl_2_. Functionalized channels were washed 3 times with this buffer prior to the experiment.

### Midplane/Cortical image and chimeric labeling

To observe PAR-3 clusters on the cell cortex, we first treated microscope coverslips with 0.1% poly-D-lysine (Sigma, P8920) and baked it until evaporated. Afterward, we added 10 µL of egg buffer and dissected the worms in the buffer. After adding 22 µm beads (white house scientific, MS0023) into the buffer, we mounted the coverslip with a glass slide and sealed with valap (a 1:1:1 mixture of petroleum jelly, lanolin, and paraffin wax). Mid-plane images were acquired on a Nikon Eclipse Ti2 microscope equipped with a 60x, 1.4 NA objective lens; a Crest XLight V3 spinning disk confocal head; and a Photometrics Prime95B camera, and controlled by MicroManager Software. 20% laser power and 200-300 ms exposure time was generally used for midplane imaging. Cortical images of mNG::PAR-3 were acquired on a Nikon Eclipse Ti2 microscope equipped with a 60x, 1.49 NA TIRF objective lens; a Gataca iLas circular TIRF illuminator; and a Photometrics Prime95B camera. 50 mW laser power, 300 ms exposure time, and a TIRF penetration depth of approximately 400 nm were generally used for cortical imaging. For the aPKC-as experiments, embryos were permeabilized with *perm-1(RNAi)* as described above, and we added 50 µM 1NA-PP1 or ethanol in SGM for both midplane and cortical imaging.

For cortical imaging with chimeric labeling, *perm-1(RNAi)* embryos were labeled with a mixture of HaloTag ligand dyes as described above. We mounted the embryos in SGM using 22 µm beads as spacers and sealed with valap. Images were acquired using the same home-built TIRF microscope described above. We used 50 mW 561 nm laser power to excite JF_585_ and 10 mW 638 nm laser power to excite JF_635_. We acquired 2000-frame movies with an exposure time of 50 ms.

### Quantification and Statistical Analysis

#### Fluorescence intensity measurements of whole-embryo images

Image intensity measurements were made using the FIJI distribution of ImageJ. To determine the RNAi knockdown efficiency of PAR-6 and PLK-1, we measured the pixel average intensity of a region of interest encompassing an entire embryo, then subtracted off-embryo background to obtain fluorescence intensity. We normalized the fluorescence intensity by dividing it by the average fluorescence intensity from non-RNAi-treated knock-in embryos.

To verify inhibition of aPKC kinase activity, we utilized the aPKC-as strain with GFP tagged PAR-2. We drew the same size of region to acquire the intensity from the anterior membrane, the posterior membrane, the cytoplasm, and the outside of the embryo. After subtracting the off-embryo background, we normalized the fluorescence intensity of anterior and posterior membranes to the intensity of the cytoplasm.

#### Data analysis for sc-SiMPull

sc-SiMPull data were processed using a published open-source software package [58]. This MATLAB package enables image registration, finding newly appearing molecules, indicating co-appearance, and analyzing photobleaching steps, and is listed in supplementary table 3. After multicolor image registration, newly appearing bait protein molecules were located. We measured the intensity of newly appearing molecules in bait and prey channels. Simultaneous signal increases in the bait and prey channels indicate co-appearing molecules, which are indicative that the two labeled proteins are in complex. In addition, we analyzed the photobleaching steps from bait and prey channels to identify the number of PAR-3 and aPKC molecules in each complexes [51].

To analyze the size distribution of PAR-3 oligomers, we plotted the mean number of observations per embryo of each cluster size. We calculated the 95% confidence interval of the mean by bootstrapping, using the built-in bootci function in MATLAB. The datapoints having too less observation, which made the 95% confidence interval overlap to 0, are marked with hollow points with no confidence interval. To assess statistical significance of observed differences between experimental groups, we calculated an effect size, expressed as log_2_ fold change, and its bootstrap 95% confidence interval. To prevent calculation errors due to division by zero or taking the logarithm of zero, we replaced zero values with 10^-10^. We designated differences between experimental groups as significant if the 95% confidence interval of the effect size did not overlap 0. To fit the PAR-3 population with exponential fits, we applied MEMLET to fit our data [101]. We used the following equation to fit the PAR-3 interacting or not interacting with aPKC:

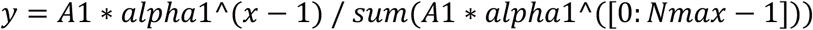

Nmax is the maximum size within the population. After fitting, we perform 1000 bootstrap samples to calculate the 95% interval of alpha1 and A1. We plot the exponential fit on the raw PAR-3 population based on A1 and alpha1.

To measure the binding affinity between aPKC and PAR-6 (Supplementary Figure 2D-E), we plotted the average co-appearance percentage over time and fit an exponential decay function to obtain k_off_ and its 95% confidence interval [58].

### Data analysis for cortical imaging with chimeric labeling

We analyzed chimeric labeling data as previously described [71]. Briefly, we used UTrack (Supplementary table 3) to identify membrane-bound Halo::PAR-3 clusters labeled with sparse JF_585_ Halo dye and measured the brightness of JF_585_ signals [102]. Afterward, we utilized a custom MATLAB package, listed in supplementary table 3, to measure the brightness of abundant JF_635_ and mNG signal colocalized with these PAR-3 clusters [71]. To convert the brightness of abundant JF_635_ to an estimated number of PAR-3 molecules, we first measured the intensity ratio of single JF_585_ and JF_635_ molecules, obtained from calibration experiments where both dyes were diluted to single-molecule levels. Then, we normalized JF_635_ intensities in experimental samples to the intensity of sparse JF_585_ in the same cluster. This approach corrects for variations in intensity due to defocus, variable eggshell thickness, and differences in Z position between clusters [71]. From the same data, we determined whether each PAR-3 cluster contained mNG::aPKC by setting a background-subtracted intensity threshold of 0 on the mNG channel. We thereby classified each cluster based on its oligomer size and PAR-3 and whether it contained or lacked aPKC. Afterward, we applied bootstrapping analysis to calculate the mean observations per embryo, effect size and 95% confidence interval, as for sc-SiMPull data. To prevent calculation errors due to division by zero or taking the logarithm of zero, we replaced zero values with 10^-10^.

## Supplementary tables

**Table S1.**
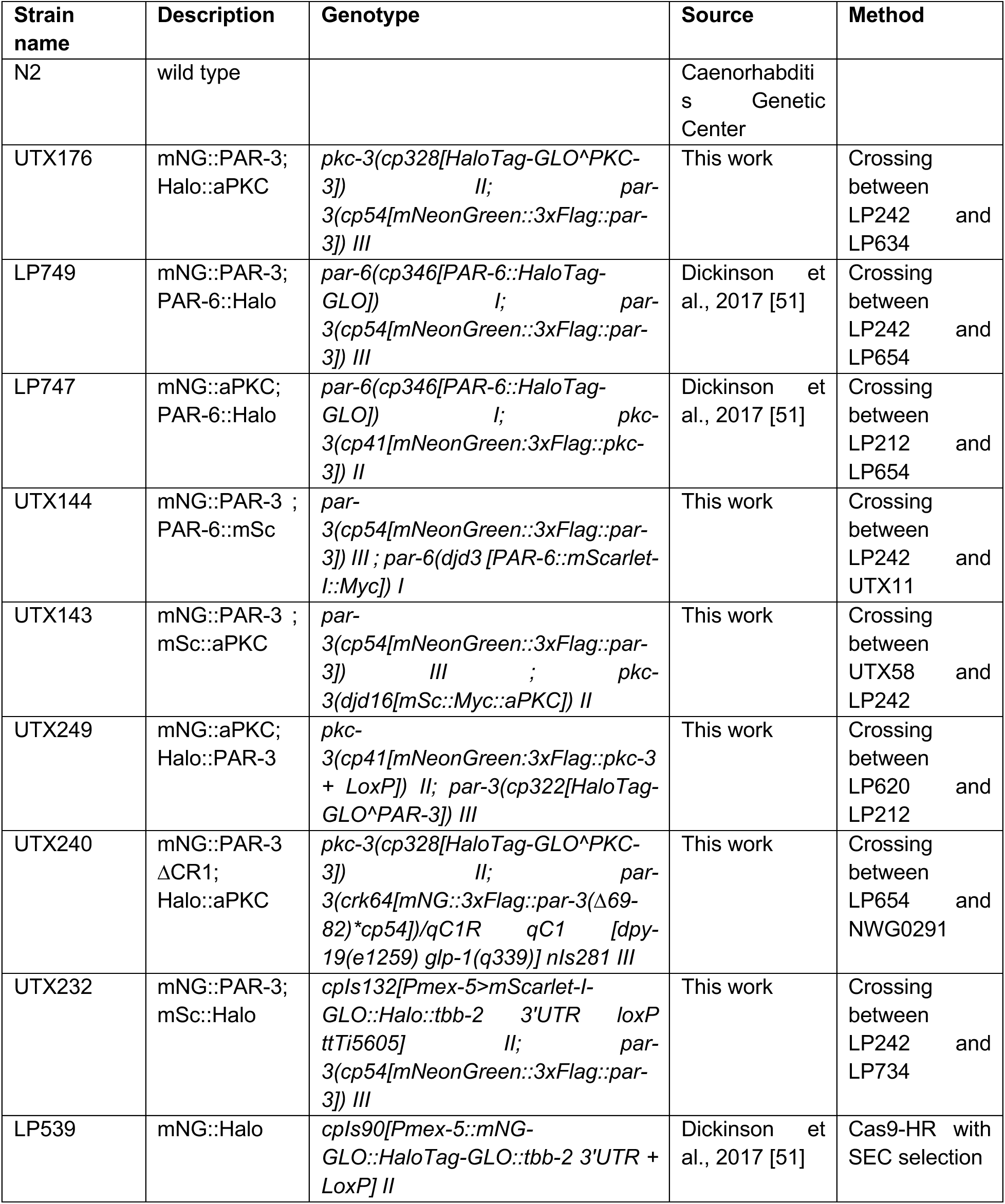
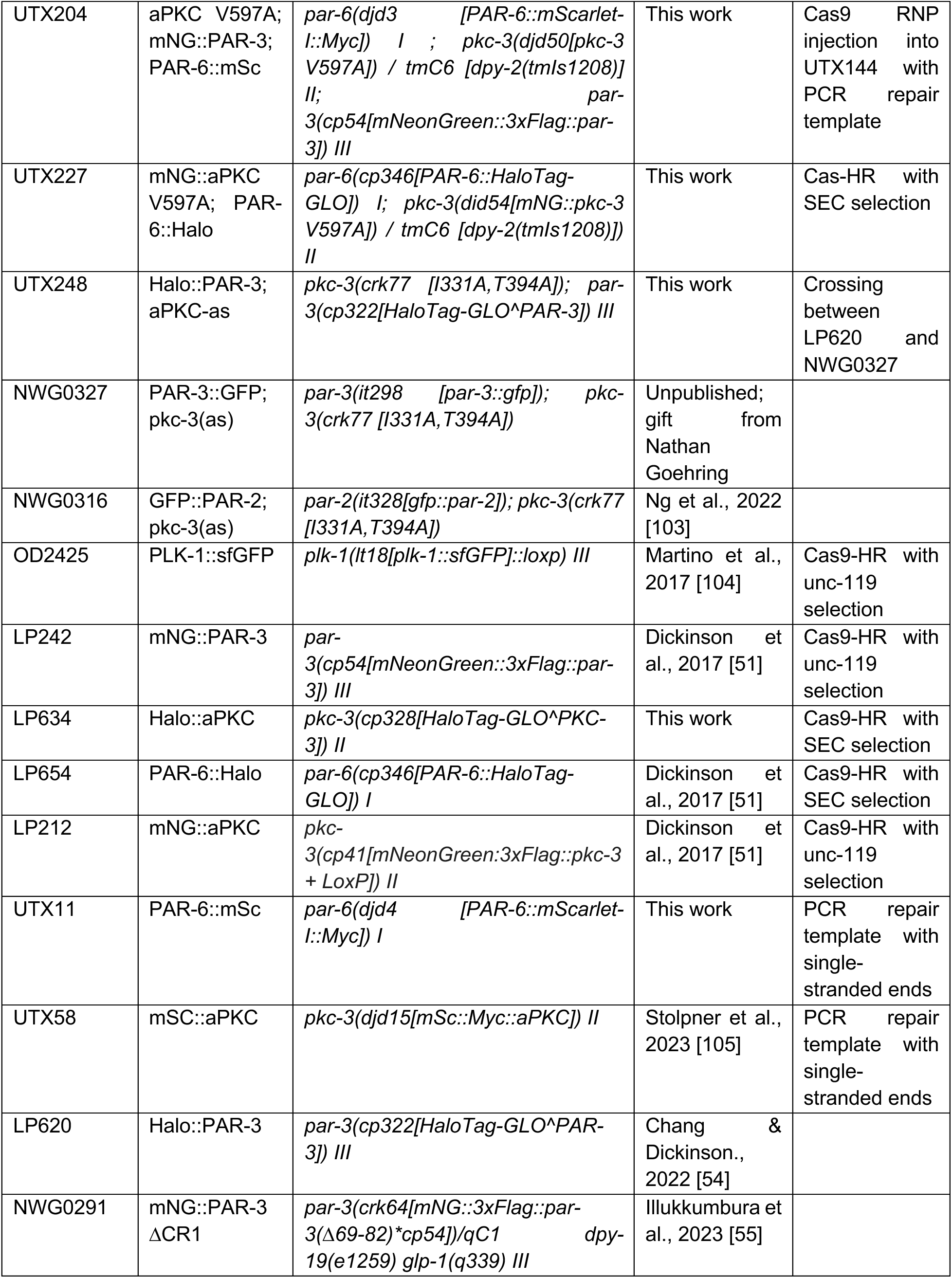
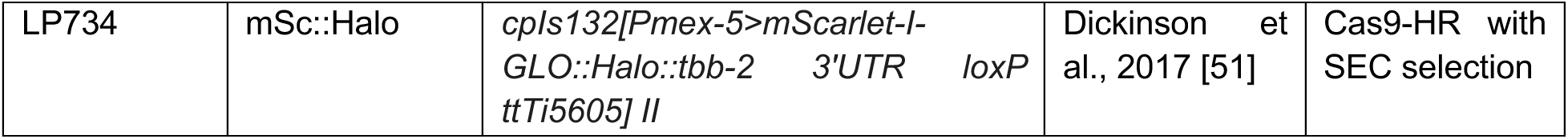
*C.elegans* strains.

**Table S2.** Key chemical reagents.

| Name | Source | catalog number |
| --- | --- | --- |
| Anti-mNeonGreen nAb | Bulldog Biosciences | NT250 |
| Biotinylated Anti-mNeonGreen nAb | Nano-tag Biotechnology | N3205 |
| 1NA-PP1 | Cayman Chemical Company | 10954 |
| ATP | New England Biolabs | P0756S |

**Table S3.** RNAi library.

| Name | Source |
| --- | --- |
| *C. elegans* feeding RNAi clones including *par-6*, *perm-1*, *plk-1* | Gifts from Jonathan Pierce; Kamath et al [106] |

**Table S4.** Software.

| Name | Reference | Purpose | Link |
| --- | --- | --- | --- |
| SiMPull acquisition software | Sarikaya & Dickinson 2021 [58] | SiMPull acquisition plugin for Micro-Manager | https://github.com/dickinson-lab/LaserLysis |
| SiMPull analysis software | Sarikaya & Dickinson 2021 [58] | analyzing single molecule imaging from SiMPull | https://github.com/dickinson-lab/SiMPull-Analysis-Software/ |
| MEMLET | Woody et al., 2016 [101] | fitting PAR-3 population with exponential distribution | https://zenodo.org/records/55586 |
| Utrack | Jaqaman et al., 2008 [102] | tracking single cortical cluster | https://github.com/DanuserLab/u-track |
| Dual labeling Analysis | Chang & Dickinson., 2022 [54, 71] | measuring intensity of HaloLigand signal from Utrack | https://github.com/dickinson-lab/DualLabeling_Analysis |
| FIJI | Schindelin et al., 2012 [107] | FIJI distribution of ImageJ | https://imagej.net/software/fiji/ |

